# The Src family kinase inhibitor drug Dasatinib and glucocorticoids display synergistic activity against tongue squamous cell carcinoma and reduce MET kinase activity

**DOI:** 10.1101/2024.10.16.618664

**Authors:** Ali N.A. Hmedat, Jessica Doondeea, Daniel Ebner, Stephan M. Feller, Marc Lewitzky

## Abstract

**Background:** Tongue squamous cell carcinoma (TSCC) is an aggressive cancer associated with a poor prognosis and limited treatment options, necessitating new drug targets to improve therapeutic outcomes. Our current work focuses on protein tyrosine kinases as well-known targets for successful cancer therapies.

**Methods:** Western blot analysis of tyrosine phosphorylation patterns in 34 TSCC lines identified Src family kinases (SFKs) as the main contributors. Inhibition of SFKs with PP2 and Dasatinib led to profound biological effects. A high-throughput screen with 1600 FDA-approved drugs was performed with three TSCC lines to discover drugs that act synergistically with Dasatinib against TSCC cell viability. Glucocorticoids emerged as potential candidates and were further investigated in 2D culture and by 3D soft agar colony formation. Dexamethasone was chosen as the major tool for our analyses. Since Dasatinib and glucocorticoids are known for their pleiotropic actions on cells, we analyzed effects on cell cycle, senescence, autophagy and cell signaling.

**Results:** A panel of 34 TSCC lines showed a surprisingly homogenous pTyr-protein pattern and a prominent 130 kDa pTyr-protein. Inhibition of SFK activity greatly reduced overall pTyr-protein levels and p130Cas tyrosine phosphorylation. It also impaired TSCC viability in 2D cell culture and 3D soft agar colony formation. A high-throughput drug combination screen with Dasatinib identified glucocorticoids as promising candidates for synergistic activity. Dasatinib and Dexamethasone combination treatment showed strong synergistic effects on Src and p130Cas phosphorylation and led to reduced p130Cas expression. Dexamethasone also suppressed phosphorylation of the MET kinase and its key substrate Gab1. On the cellular level, Dasatinib combination with glucocorticoids led to G1 cell cycle arrest, increased senescence and enhanced autophagy. This was also reflected by effects on cell cycle regulatory proteins, including CDKs and cyclins.

**Conclusion:** This work is the first to show a strong synergistic activity of Dasatinib in combination with clinically used glucocorticoids in solid tumors. Furthermore, the tyrosine kinase MET and its effector protein Gab1 are newly identified glucocorticoid targets. Given the extensive research on MET as a drug target in various cancers, our findings have the potential to advance future cancer treatments.

## Introduction

Tongue squamous cell carcinoma (TSCC) is a serious public health problem with significant morbidity and mortality [1]. It accounts for 40 to 50% of oral squamous cell carcinomas (OSCCs) worldwide, making it the most prevalent intraoral cancer [2]. Smoking, which leads to cancer-causing mutations, is clearly the major overall cause of TSCC and the combination of tobacco consumption by smoking or chewing with high alcohol consumption synergistically increases the carcinogenic effects [3]. Human papilloma virus (HPV) infections can also contribute to TSCC development [4]. Fortunately, effective immunizations against multiple major HPV strains are now available, although these vaccines are so far much less utilized than they could be. It is possible that the increased incidence of tongue carcinomas in women and younger adults is at least partially related to changes in smoking, drinking and other lifestyle habits [5].

Previous analyses, which investigated the mutational features of head and neck squamous cell carcinomas (HNSCC), including different TSCC, revealed various mutations in genes of therapeutic and prognostic significance [6, 7]. In these studies, apart from the mutations in the prominent tumor suppressor gene p53, affected pathways include EGF family receptor signaling, the mitogen-activated protein kinase (MAPK) and NOTCH signaling pathways, as well as adherens junction and focal adhesion pathways. However, many other mutations and dysfunctional regulatory mechanisms likely remain to be discovered. Unfortunately, such molecular insights have had relatively little impact on the clinical treatment outcomes for TSCC so far.

Surgery is often the primary treatment of early (stage T1 and T2) tumors of the anterior two-thirds of the tongue, in combination with radiotherapy, also for larger, more posteriorly placed lesion [8]. Total glossectomy is performed for advanced carcinomas of the tongue [9]. This is associated with a high level of morbidity [10]. Conventional chemotherapy has a limited role in the primary management of carcinoma of the tongue, but is sometimes considered as an adjunct treatment in advanced disease, like unresectable lesions or distant metastasis. The five-year survival rates for TSCC have remained essentially unchanged over the past 30 years, despite some advances in TSCC diagnosis and management [11].

Molecularly targeted therapies currently play a limited role in head and neck cancers (HNCs). Treatment with the monoclonal anti-EGFR antibody Cetuximab has shown some clinical benefits. However, although there are now numerous additional targeted therapies for patients with HNC in clinical testing [12], there is no particularly effective molecularly targeted therapy for TSCC available until now. Due to the increase in TSCC incidence, the high risk of recurrence and the poor prognosis, the search for effective clinical treatments for TSCC has become an increasingly important topic. Identification of new molecular targets may enable significantly improved treatments.

Protein tyrosine kinases (PTKs) play key roles in cellular signal transduction, cell cycle regulation, cell division, and cell differentiation. The signaling pathways regulated by protein kinases are frequent targets of mutations, leading to many human cancers [13]. Dysregulation of PTK-activated pathways, often by overexpression, gene amplifications, or genetic mutations, are causal factors underlying numerous cancer developments, as well as disease progression and drug resistance [14]. The inhibitions of kinases with drugs have produced substantial antitumor effects for some cancer types.

To detect tyrosine kinases which might become therapeutic targets in TSCC treatment we have studied tyrosine phosphorylation in a panel of 34 TSCC lines. Western blot analysis of their protein tyrosine phosphorylation status, using a well-established anti-phosphotyrosine antibody, revealed a surprisingly homogenous pTyr-protein pattern throughout the TSCC panel with a prominent 130 kDa pTyr-protein. This led us to focus initially on the potential involvement of the multi-site docking protein p130Cas (BCAR1) and Src family kinases (SFK) in TSCC. Inhibition of SFK activity with two inhibitors, the tool compound PP2 [15] and the clinically used drug Dasatinib (DAS) [16], greatly reduced overall pTyr-protein levels and p130Cas tyrosine phosphorylation on specific epitopes. Moreover, TSCC viability in standard cell culture and 3D soft agar colony formation was greatly diminished.

Since monotherapies with anti-cancer drugs are frequently ineffective or short lasting in their activity and thus require additional drugs to accomplish useful clinical responses [17-19], we performed a high-throughput screen with 1600 FDA-approved drugs, to discover candidate molecules that achieve synergistic activity against TSCC lines when combined with DAS. This screen identified glucocorticoids (GCs), including Dexamethasone and Fluticasone, as a promising group of drugs for further analyses. GCs have been widely used in medicine for decades, including as part of some cancer treatment regimens [20, 21]. The expansion of their clinical use in cancer therapies could be swift and cost-effective, with only moderate concerns about off-target toxicities. After more detailed studies, GCs were found to induce rapid and prolonged inhibition of the MET tyrosine kinase and its key substrate protein Gab1. This suggests that GCs may have a useful role to play in the clinical inhibition of MET signaling in tumors.

## Results

### Similarity of tyrosine phosphorylation patterns and prominence of a 130 kDa pTyr-protein in a panel of 34 TSCC lines

Tyrosine kinases are frequent drivers of human tumor development but remain only marginally studied in TSCC. Identifying tyrosine kinases that are overexpressed or hyperactivated in TSCC should enable us to think more rationally about targets to counteract this devastating disease. To gain a first insight into the abundance and sizes of pTyr-proteins in TSCC, total cell lysates from 34 cell lines (30 established from primary tumors, 4 from metastases) were subjected to Western blot analysis of the protein tyrosine phosphorylation status using an anti-phosphotyrosine antibody (**Figures 1A and S1**). Further information on the cell lines can be found in **Suppl. Table 1** and in [22].

**Figure 1.**
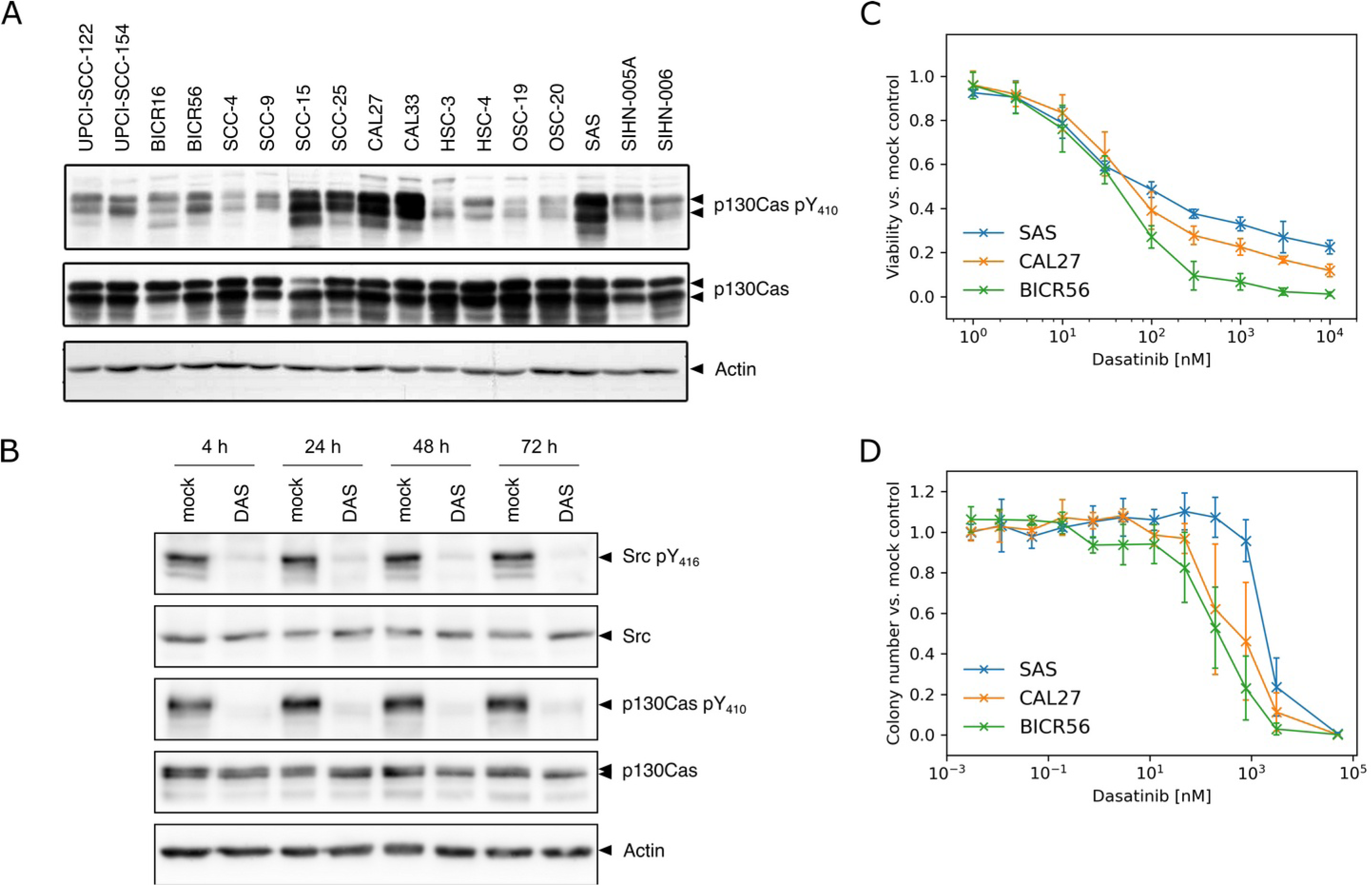
Dasatinib reduces the growth of TSCC cells and causes inhibition of Src pTyr416 and p130Cas pTyr410 phosphorylation. (A) Total TSCC cell lysates were separated by SDS-PAGE and analyzed by Western blotting with the indicated antibodies. Actin serves as loading control. (B) Western blot analysis of the total level and the phosphorylation level of Src and p130Cas in SAS cells treated with DMSO as vehicle control (mock) or 100 nM of Dasatinib (DAS) for the indicated time points. (C) Dose-response curves for DAS in TSCC lines as determined by resazurin assay after 72 h treatments. Values are normalized to mock-treated cells. The dose-response curves show mean values ± SD for three independent experiments with six parallel measurements each. (D) Dose-response curves of TSCC lines treated with DAS as determined by colony formation in 3D soft agar assay. Cells were treated with DMSO as vehicle control (mock) or different concentrations of DAS. Colony numbers were normalized to mock-treated cells. The dose-response curves show mean values ± SD of three independent experiments with at least 2 wells counted in each case.

Cells were fed with an excess of culture medium 48 h before harvesting, to minimize medium effects on pTyr-protein and to obtain information on the cells’ ‘steady state’ phosphorylation. Unexpectedly, all TSCCs exhibit fairly similar tyrosine phosphorylation patterns with particularly notable bands at around 130 kDa, possibly indicative of a prominent role of a single kinase or kinase family in TSCC. This is in stark contrast to the diverse pTyr patterns seen in other cancer cell line panels, for example a panel derived from colorectal carcinomas, which display a much greater variability concerning pTyr-protein abundances and band patterns [23]. p130Cas is a well-known and major substrate protein of several oncogenic tyrosine kinases, including Src and Abl family kinases [24-26] and was therefore analyzed in the TSCC panel.

### Identification of p130Cas (BCAR1) as ubiquitous pTyr-protein in the TSCC line panel

p130Cas contains numerous potential tyrosine phosphorylation sites that are mostly clustered in the ‘substrate region’ of the protein. p130Cas (later also named BCAR1) has been characterized as a docking and signal processing platform for multiple signal transduction proteins [27-29]. To check for the total protein levels and phosphorylation status of p130Cas, total cell lysates from the 34 TSCC lines were analyzed by Western blotting (**Figures 1A and S1**).

The p130Cas protein is expressed throughout all cells of the cell line panel. The protein is also phosphorylated in all TSCC lines on Tyr 410 (Y410), a major phosphorylation site and binding motif for Crk family adaptor proteins [30]. However, since p130Cas is well known to be phosphorylatable on more than a dozen tyrosines (see phosphosite.org and references therein), this phospho-antibody, raised against a single phosphorylation site of p130Cas, does not necessarily reflect the overall phosphorylation detected by the pTyr-antibody. From these initial results, we surmised that p130Cas is probably a major component of the 130 kDa phospho-tyrosine band detected in all TSCC lines of the panel and that it might be a potential therapeutic target for tongue cancer. However, the non-catalytic nature of p130Cas makes it difficult to develop specific inhibitors. Therefore, alternative strategies are needed to inhibit p130Cas activity.

### Src family kinases are essential for maintaining the tyrosine phosphorylation of p130Cas and other pTyr-proteins in TSCC lines

Previous work has shown that the C-terminal region of p130Cas contains a Src kinase binding domain and that the association between p130Cas and activated Src kinase is essential for the tyrosine phosphorylation and activation of p130Cas by Src at least in some cells [31]. Therefore, SFK inhibitors have been proposed as useful tools to inhibit p130Cas activity. Hence, the effect of the SFK inhibitor compound PP2 [32-34] on the overall pTyr-protein levels and p130Cas activity was analyzed in SAS cells (**Figure S2A**). SAS cells were chosen for these initial experiments since they are widely studied, fast growing and highly motile, presumably representing a particularly aggressive form of TSCC.

Upon PP2 treatment of SAS cells, a substantial drop in tyrosine phosphorylation was observed at all time points. A control compound, PP3, which lacks SFK blocking activity, did not lead to detectable pTyr pattern changes. p130Cas pY410 phosphorylation was also clearly reduced after PP2 application. During these experiments, striking effects on SAS cell survival and proliferation were noticed, which led us to investigate the crucial proliferation-regulating MAP kinases Erk1 and Erk2 as well. Severe reductions in the levels of active Erk1/2 were apparent. Blotting with a Src pY416 antibody, expectedly, also showed a rapid decline in the activity-regulating tyrosine phosphorylation of SFK. This tyrosine residue is well-known to contribute to the regulation of Src activity [35] and the Tyr416 phospho-antibody recognizes several SFK members that share a high degree of homology in their relevant protein epitopes [23].

### Live cell imaging of SAS cells reveals pleiotropic effects of tyrosine kinase inhibition with PP2

SAS cells treated with PP2 show a striking growth inhibition compared to DMSO-treated control cells. To analyze this further, live cell imaging was performed. Detailed inspection of the videos of SAS cells treated with DMSO or PP2 further documented that PP2 can exert several biological effects (Video S1). Not only does PP2 inhibit the swarm-like cell migration of SAS cells and their cell proliferation, but it also induces a substantial change in cell morphology, and, possibly, a reversion of epithelial-mesenchymal transition (EMT) [34]. Before PP2 application, individual cell boundaries are clearly visible. These disappear upon cell exposure to PP2, but not DMSO alone, yielding more dense epithelial sheets. Starting after approximately 24 h of treatment and more prominently visible from ca. 40 h onward, membrane blebbing and cell death is also observed. All of these effects of tyrosine kinase inhibition would be expected to be anti-oncogenic.

### Dasatinib reduces TSCC lines growth in a dose-dependent manner and inhibits Src family kinase activity

Since PP2 is just a tool compound and not approved for cancer treatments, we decided to further investigate the SFK inhibition effect on the reduction of p130Cas activity and the growth of TSCC using the clinically important SFK inhibitor Dasatinib (DAS) [36]. DAS is approved, for example, for the treatment of chronic myelogenous leukemia (CML) refractory to or intolerant of Imatinib, but also as first-line treatment of chronic-phase CML and for patients with Philadelphia chromosome-positive acute lymphoblastic leukemias [36].

Initially, SAS cells were treated with 100 nM DAS, a clinically achievable concentration [37], for increasing durations (from 4 h up to 72 h). DAS caused an almost complete inhibition of SFK and p130Cas phosphorylations, as determined by western blots with anti-pY416 Src and anti-pY410 p130Cas, respectively (**Figure 1B**). This inhibition was apparent at all time points.

It should be noted that DAS inhibits not only Src but also Abl family kinases, which could contribute to the suppression of overall pTyr levels in the TSCC lines. The reduced p130Cas phosphorylation may also be the result of Abl inhibition since p130Cas is also well-known Abl substrate [38, 39]. To determine if SFK inhibition accounts for the observed reduction of overall pTyr levels and p130 Cas pY410, we also tested the clinically used Abl family kinase inhibitor Imatinib on SAS cells. In addition to Abl family kinases, Imatinib can also inhibit c-Kit, PDGFR and other kinases, but it has no prominent effect on SFKs [40]. DAS, but not Imatinib, reduced pTyr, p130Cas pY410 and SFK pY416 levels in a dose-dependent manner (**Figure S2B**). These data indicate that SFK inhibition by DAS and not Abl inhibition is very likely responsible for massively decreasing pTyr-protein levels in TSCC.

Subsequently, three TSCC lines (SAS, CAL27 and BICR56) were selected from the large TSCC panel to represent different levels of tyrosine phosphorylation (**Figure S1**) and for their ease of global availability. These cell lines were treated with different concentrations of DAS for 72 h. Cell viability under standard cell culture (2D) conditions was then determined by resazurin assay. As shown in **Figure 1C**, cell viability was reduced in a dose-dependent manner in all three cell lines. These results indicate that DAS has clear activity against TSCC cells in 2D culture. The SAS cell line shows a lesser response to DAS compared to the other 2 cell lines, with BICR56 being the most sensitive cell line. DAS did not eliminate all SAS and CAL27 cells, even at very high concentrations (10 µM) after 72 h of exposure, which suggests that the effects of DAS on cell growth are, at least in part, antiproliferative/cytostatic and not entirely cytotoxic.

To investigate the effects of DAS on TSCC cell lines in 3D culture, a classical soft agar assay, which can, to some degree, mimic the growth of cancer cells in actual tissues, was employed. TSCC cells were seeded into soft agar layers and treated with different concentrations of DAS. Cells were maintained in culture for 14 days to allow the formation of colonies. DAS greatly reduces colony formation SAS, CAL27 and BICR56 in a dose-dependent manner, as shown in **Figures 1D and S4C**.

### A high-throughput drug combination screen identifies drugs acting synergistically with Dasatinib on TSCC lines

Small, early phase clinical trials with exploratory SFK inhibitor treatments of unselected head and neck cancer patients have so far failed to show significant clinical benefits [17-19]. The underlying reasons for this lack of clinical activity as monotherapies remain to be identified. However, nearly all effective cancer treatments are currently based on the additive or even synergistic actions of several drugs applied in combination [41, 42].

To identify drug candidates that display synergistic activity together with DAS from the large collection of existing FDA-approved drugs, we performed high-throughput screens using the Pharmakon 1600 library on SAS, CAL27 and BICR56 cells. Details of the screens are described in Materials and Methods. Briefly, drugs that showed an outstanding ratio of cell metabolic activity impairment in combination with DAS compared to cell metabolic activity impairment for the drug alone, in more than one cell line, were considered for further analyses (**Supplementary Data S1).** For the SAS cell line, a majority of candidate drugs were effectors of glucocorticoid or β-adrenergic signaling stress receptor signaling pathways. Two drugs, the glucocorticoid Fluticasone and β2-adrenergic receptor agonist Isoproterenol were also found to have synergistic activity on the BICR56 cell line. Initial experiments with β2-adrenergic receptor agonists were not successful and so it was decided to focus on the potential synergy of DAS and glucocorticoids for further work.

To assess the possible synergistic activity between DAS and Fluticasone (FL), SAS cells were initially treated with increasing concentrations of FL as single agents, or with a fixed concentration of FL in combination with increasing concentrations of DAS. Cell viability was examined by resazurin assay after 72 h of treatment. FL alone has no significant effect on SAS cell viability up to a concentration of 10 µM (**Figure S3A**). Next, clinically relevant concentrations up to 100 nM of FL [43] were tested in combination with DAS. 25 nM of FL was sufficient to significantly enhance the cell toxicity of DAS in the SAS cells (**Figure S3B**).

To confirm the cytotoxic effect of the DAS+FL against TSCC, trypan blue dye exclusion assays were performed. 50 nM of DAS was combined with 25 nM of FL. After 72 h treatments, cells were trypsinized, mixed with dye and the numbers of viable cells were counted with an automated cell counter and normalized to mock-treated cells (**Figure S3C**). The proliferation of all TSCC lines tested was strongly reduced in the presence of DAS+FL versus DAS alone. Again, individual application of FL showed no significant effect. The trypan blue assay results thus confirm the synergistic activity of the DAS+FL combination treatment found in the resazurin assay.

While these results were certainly encouraging, it should be noted, that FL suffers from low oral bioavailability and its clinical utilization is limited to topical routes of administration [44]. Therefore, it was decided to investigate systemically applicable GCs, to determine whether they also show synergistic effects in TSCC lines in combination with DAS.

### Dasatinib + Dexamethasone combination treatment shows a strong synergistic effect against TSCC cells in 2D and 3D cultures

Dexamethasone (DX) is a long-acting GC, available in multiple dosage forms, which is used to treat a large variety of diseases [45]. It therefore seemed to be a highly suitable candidate in search for synergistic GC - DAS activities. SAS cells were thus treated with DX or DAS as single agent or with DAS+DX combinations and the cell viability was measured by resazurin assay after 72 h. DX treatment alone showed no substantial effect on the metabolic activity of SAS cells at concentrations of up to 10 µM (**Figure 2A**). 100 nM of DX sensitized SAS cells to DAS and synergistically enhanced its effect on cell viability (**Figure 2B**). All investigated TSCC lines showed clear responses, with the effect being more pronounced for the SAS cell line than for CAL27 and BICR56 (**Figures S4A and S4B**). To further examine this synergistic interaction in 2D culture, trypan blue dye exclusion assays were conducted to determine the number of viable cells after each treatment. 50 nM of DAS was combined with 100 nM of DX. After 72 h, viable cells were counted and normalized to mock-treated cells. A combination of DX and DAS reduced the number of viable cells drastically and more strongly than DAS alone (**Figure 2E**).

**Figure 2.**
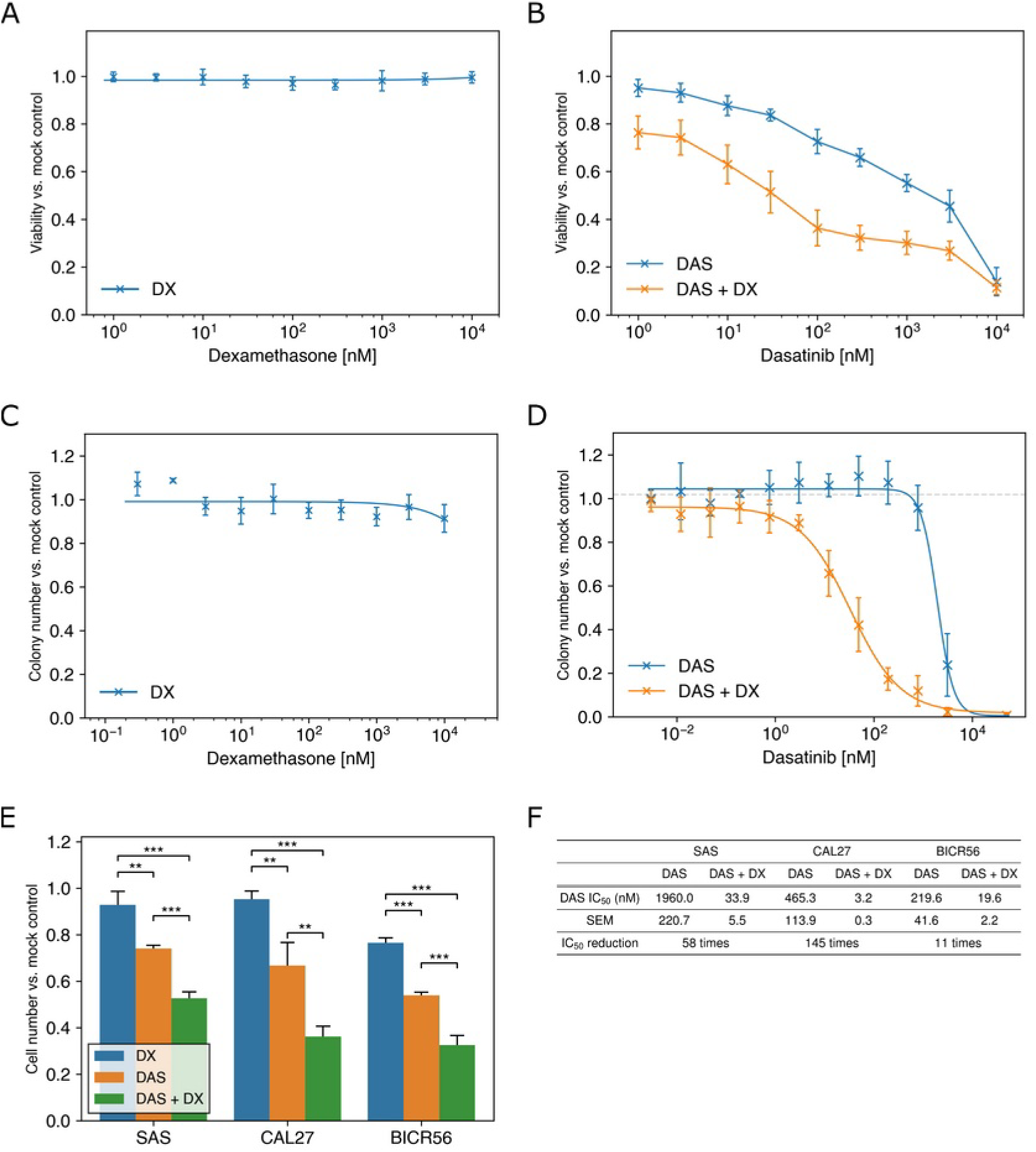
Dasatinib + Dexamethasone combination treatment shows a synergistic effect on TSCC cells in 2D and 3D cell cultures. (A) Dose-response curves of SAS cells treated with different concentrations of DX in 2D culture for 72 h as analyzed by resazurin assay. (B) Dose-response curves of SAS cells treated with different concentrations of DAS with or without 100 nM DX for 72 h. Cell viability was measured by resazurin assay. (C) Colony numbers in 3D soft agar culture for SAS cells treated with different concentrations of DX. (D) Dose-response curves of SAS cells treated with DAS with or without 100 nM DX, as determined by colony numbers in 3D soft agar culture. (E) TSCC lines were treated with DMSO as vehicle control (mock), 50 nM of DAS, 100 nM of DX, or DAS+DX for 72 h. Cell numbers were evaluated by staining cells with trypan blue and counting of viable cells with an automated cell counter. Asterisks indicate statistical significance for T-tests with p-values at less than 0.05 (*), less than 0.01 (**) or less than 0.001 (***), respectively. In A to E, data were normalized to mock-treated cells and values are expressed as means and standard deviation obtained from three independent experiments. (F) Comparison of the average IC_50_ values for DAS alone and DAS+100 nM DX in TSCC lines assessed by 3D soft agar culture. IC_50_ values were derived by fitting a four-parameter log-logistic function. Values are presented as the means ± SEM for three independent experiments.

Subsequently, soft agar colony formation assays were performed to investigate the effects of DAS+DX treatment on anchorage independent growth. As shown in **Figure 2C**, DX by itself did not cause any substantial colony formation reduction in SAS cells, even at a very high dose of 10 μM. In contrast, DAS on its own reduced colony formation in a dose-dependent manner (**Figure 2D, Figure S4C**). This was also observed in Cal27 and BICR56 cells (**Figures S4D and S4E)**. The dose-response curves for the combination treatments show that 100 nM of DX and DAS synergistically reduce colony formation in TSCC lines. SAS cells, which showed the weakest response to DAS treatment alone, with an IC_50_ value of around 2000 nM, displayed high sensitivity to the combination treatment. The IC_50_ value was reduced by a factor of 58 when adding 100 nM DX (**Figure 2F**). CAL27 cells displayed an even greater responsiveness to the DAS+DX treatment. The IC_50_ decreased by a factor of 145 from around 500 nM with DAS alone to 3.2 nM with DX+DAS. DAS+DX treatment showed less synergistic activity in BICR56 cells, but the IC_50_ was still reduced 11 times by the combination treatment.

### Dasatinib + Dexamethasone combination treatment leads to a pronounced reduction of p130Cas expression

Next, we investigated the molecular effects of DAS in combination with DX on Src and p130Cas. TSCC lines were treated with DMSO (mock), 50 nM of DAS, 100 nM of DX or 50 nM DAS plus 100 nM DX for 72 h, followed by Western blot analyses. As before, DAS alone strongly inhibited Src and p130Cas phosphorylation. This was not enhanced by addition of DX (**Figure 3**). However, DAS+DX reduced the total levels of p130Cas. The same was also observed for DAS+FL treatment (data not shown). This suggests that the biological effects seen with DX could, at least to some degree, result from effects on the p130Cas protein. However, GCs are well-known to induce rather pleiotropic effects on cell signaling in different cell contexts [46], so various other signaling pathways have to be considered.

**Figure 3.**
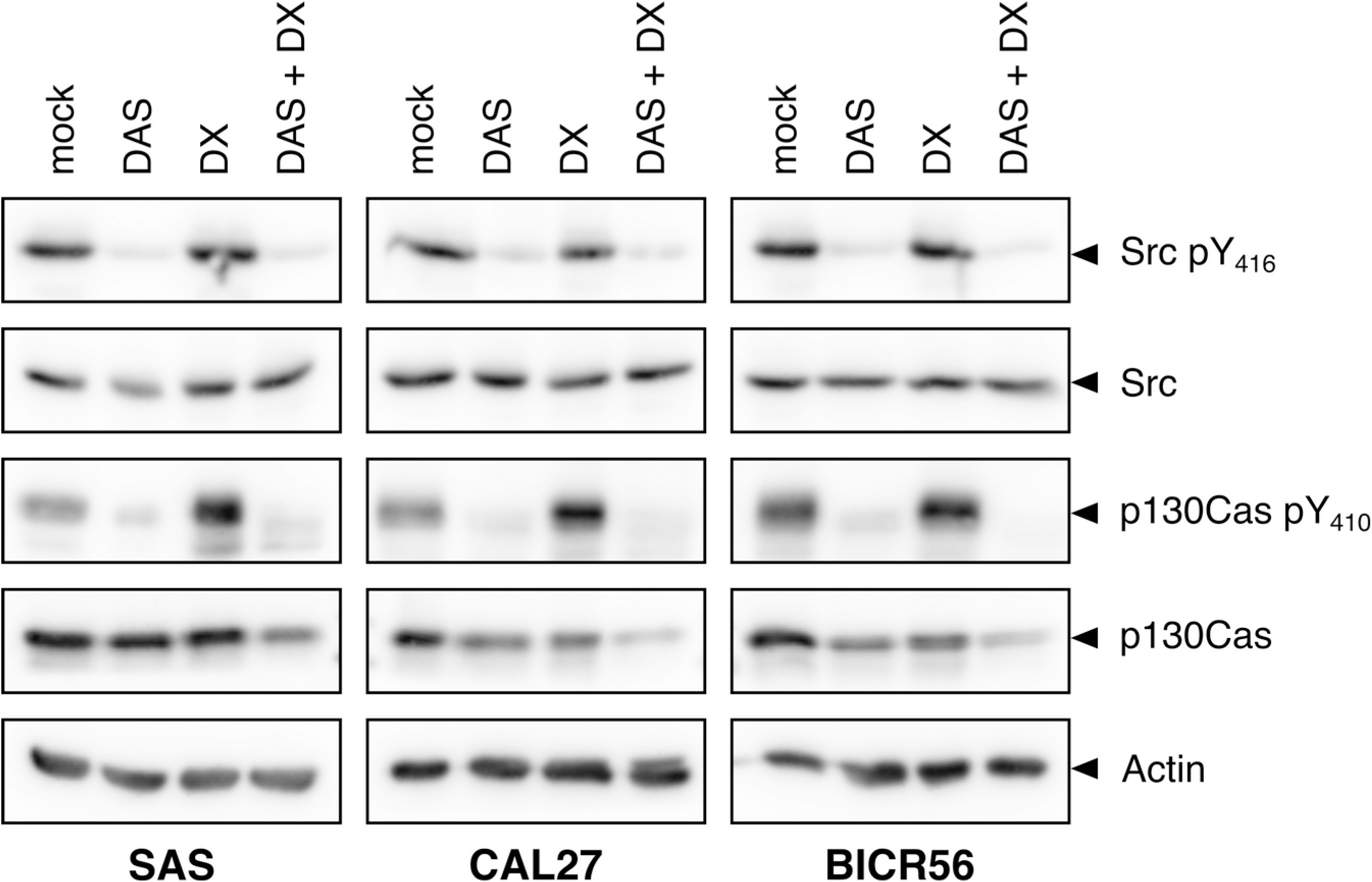
Dasatinib inhibits phosphorylation of Src and p130Cas, Dasatinib + Dexamethasone co-treatment reduces p130Cas protein levels. Western blot analysis of Src and p130Cas expression, as well as Src and p130Cas phosphorylation, in SAS, CAL27 and BICR56 cells treated with DMSO (mock control), 50 nM DAS, 100 nM DX, or a combination of DAS+DX for 72 h.

### Dexamethasone treatment suppresses MET kinase signaling in TSCC lines

Src has been shown to interact with a range of receptor tyrosine kinases (RTKs), either as a downstream target, or even as an upstream effector, as we have previously shown for the RTK mesenchymal-epithelial transition factor (MET) in colorectal cancer cell lines [23]. Overexpression of MET has been observed in more than 80% of head and neck squamous cell cancers and preclinical and clinical studies have linked MET overexpression to cellular proliferation, invasion, migration, and poor prognosis [47]. Thus, we next analyzed the expression and phosphorylation of MET and its key substrate protein Gab1 upon treatment with DAS and DX, either alone or in combination for 72 h, in the three TSCC lines.

MET phosphorylation at Y1234/1235, Gab1 phosphorylation at Y627 (a docking site for the proliferation-driving phosphatase Shp2) and total protein levels were assessed by Western blotting. As shown in **Figure 4A**, MET phosphorylation was somewhat reduced by DAS, but, surprisingly, greatly diminished upon DX treatment. The same pattern was observed for Gab1 pY627. Both drugs showed no major effect on the total MET and Gab protein levels, although the mature form of the MET receptor kinase (upper band) may be somewhat reduced, possibly indicating some effect on proteolytic MET processing.

**Figure 4.**
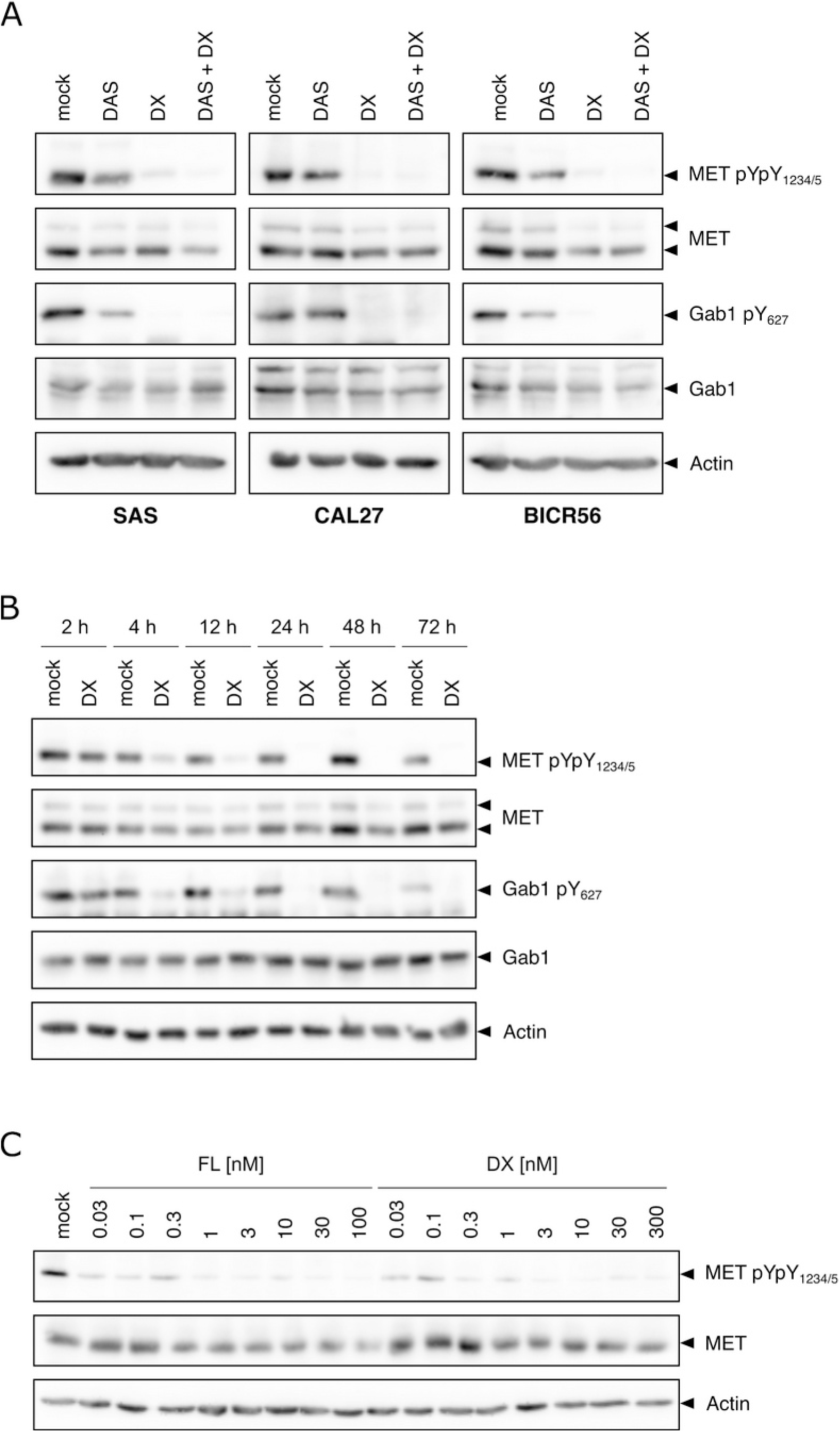
Dasatinib + Dexamethasone combination treatment suppresses MET kinase signaling. (A) Western blot analysis of MET and Gab1 expression and phosphorylation in TSCC cells treated with DMSO (mock), 50 nM of DAS, 100 nM of DX, or a combination of DAS+DX for 72 h. Antibodies that detect the phosphorylation of MET at Tyr1234/1235 and of Gab1 at Tyr627 were used to monitor changes in phosphorylation levels. (B) Western blot analysis of MET phosphorylation in SAS cells treated with DMSO (mock) or 100 nM of DX for the indicated time points. (C) Western blot analysis of MET phosphorylation in SAS cells treated with the indicated concentrations of FL or DX for 72 h.

To determine how fast the DX effects on MET and Gab1 phosphorylation occur, we incubated SAS cells with DX for up to 72 h and collected samples at different time points. As shown in **Figure 4B**, MET dephosphorylation became obvious after 4 h, reached a maximum at 24 h and persisted for at least 72 h. This correlated with the repression of phosphorylation at Y627 of the MET downstream target Gab1. The somewhat delayed inhibition of MET phosphorylation suggested to us that DX is probably not a direct inhibitor of MET kinase activity. To test this assumption, a MET kinase activity assay was performed.

Briefly, MET was immunoprecipitated from SAS cell lysates and incubated with ATP plus a recombinantly expressed fragment of Gab1 (aa 613-694) as kinase substrate. DMSO, 100 nM DX, 500 nM DX or 100 nM Crizotinib (a potent and direct MET kinase inhibitor) were added to the kinase reaction. The results, shown in **Figure S5**, indicate that, unlike Crizotinib, DX is no direct inhibitor of MET kinase activity at a concentration of 500 nM, which is approximately 5 times higher than the pharmacologically achievable plasma concentration.

To define the potency of DX and to investigate, if the inhibitory effect on MET can also be seen with another GC, SAS cells were treated with different concentrations of DX or FL for 72 h in the absence of any DAS. Strikingly, subnanomolar concentrations of either GC were already sufficient to inhibit MET phosphorylation substantially (**Figure 4C**). The exact mechanism of the surprising and potent effect of GC on the MET - Gab1 signaling pathway remains to be further investigated. In addition, GCs, as mentioned above, may have significant effects on additional cellular signaling pathways.

### Dasatinib + Dexamethasone combination treatment induces G1 cell cycle arrest and senescence in TSCC cells

In a subsequent set of experiments, we aimed to determine the effects of DAS, DX and their combination on the cell cycle and its regulatory proteins. The three TSCC lines were treated with DMSO (mock), 50 nM DAS, 100 nM DX, or a combination of DAS+DX for 72 h and thereafter stained with DAPI for determination of their DNA content by flow cytometry (**Figure 5A**). Individually, both compounds induced a slight increase of cells in the G1 phase, while combination treatment caused more significant G1 arrests (SAS: mock 59.7% ± 1.9% vs. DAS+DX 86.2% ± 0.8%, n = 3). As expected, cells in S phase were notably reduced (SAS: mock 29.2% ± 0.1% vs. DAS+DX 7.6% ± 0.1%, n = 3), as were cells in G2/M phase (SAS: mock 11.1% ± 0.3% vs. DAS+DX 6.2% ± 0.1%, n = 3). No significant sub-G1 peak increases, indicative of substantially elevated cell death, were found (data not shown).

**Figure 5.**
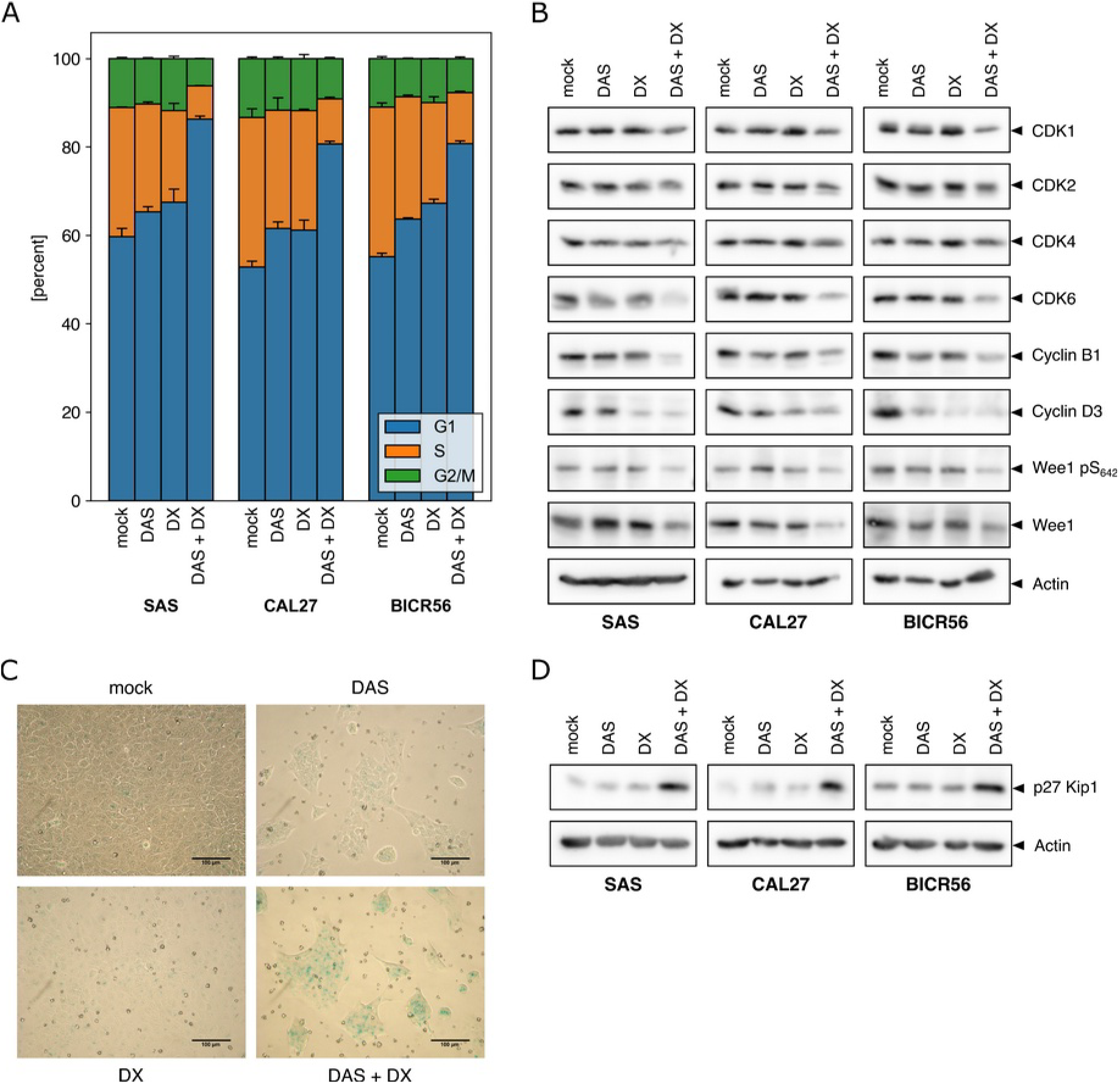
Dasatinib + Dexamethasone combination treatment induces cell cycle arrest in G1 and senescence. Cells were treated with DMSO (mock), 50 nM of DAS, 100 nM of DX or a combination of DAS+DX for 72 h. (A) Treated TSCC lines were stained with DAPI and DNA contents were determined by flow cytometry. 10,000 cells were analyzed for each experiment. Cell populations are displayed as percentage of cells at each cell cycle phase relative to the total population. Results are expressed as means and standard deviation obtained from 3 independent experiments. (B) Western blot analyses of CDKs, cyclins and Wee1 as indicated. (C) Representative pictures of SAS monolayer cultures stained with senescence-associated-β-Galactosidase (SA-β-Gal). SA-β Gal positive staining (blue color) indicates aged cells. The scale bar length indicates a length of 100 µm. (D) Western blot analysis of p27Kip1 protein expression.

Next, the protein expression levels of several crucial cell cycle regulators were examined in SAS, CAL27 and BICR56 cells using the same treatment conditions (**Figure 5B**). Western blot analyses showed that the levels of the cyclin-dependent kinases CDK1, possibly CDK4 and, especially, CDK6 were reduced upon DX+DAS co-treatment. In addition, we investigated the expression of Cyclin B1, together with CDK1, a gatekeeper for the entry into mitosis, and Cyclin D3 which interacts with CDK4 and CDK6 to regulate G1/S-transition. Both cyclins were reduced by DAS+DX treatment. This agrees with the observed changes in cell cycle phase distributions (**Figure 5A)**. An additional key player of cell cycle regulation, the Wee1 kinase, an inhibitor of mitosis entry [48], was also examined. The observed total reduction in Wee1, together with dephosphorylation of pSer642 upon treatment with DAS+DX (**Figure 5B**), is expected to reduce nuclear Wee1 and increase its cytoplasmic localization [49]. The changes in CDKs, Cyclins and Wee1 levels, observed so far, warrant a more extended and detailed future investigation of cell cycle regulator signaling in order to determine the functionally relevant alterations elicited by DAS+DX combination treatment.

Besides cell cycle changes, glucocorticoids have also been reported to affect other cell fates, including cellular senescence [50, 51]. These effects can vary, depending on cell type, length of exposure and physiological context [52-54]. Therefore, the consequences of DAS+DX treatment in relation to cellular senescence are hard to predict. To gain a first insight into this, senescence-associated beta-galactosidase (SA-β-gal) assays were performed as described in the methods section. SA-β-gal positive blue staining, which indicates senescent cells, was detected by light microscopy (**Figure 5C and S6A+B**). Co-application of DAS and DX for 72 h increased the blue staining. In the following, the expression levels of p27Kip1 were evaluated. p27Kip1 has been reported as a driver for DX-induced senescence and is also a more general inducer of cell cycle blockades due to its interactions with CDK/Cyclin complexes in various normal and cancer cells [52, 55] (**Figure 5D**). Not unexpectedly, p27Kip1 was upregulated in all three TSCC lines after co-treatment with DAS+DX.

### Dasatinib + Dexamethasone treatment induces autophagy in TSCC cells

Another biological response altered in cancer cells by some drugs is autophagy, the degradation of cellular material by lysosomes [56]. GCs have been shown to induce autophagy in lymphoid leukemia [57] and pancreatic ductal adenocarcinoma cells [58]. Autophagy has context-dependent roles in cancer. It is thought that autophagy can prevent the development of some cancers, but, once established, it may help cancer cells to survive under supply stress conditions [59-61].

To examine if DAS alone, DX alone, or the combination of DAS plus DX impact on autophagy in SAS, CAL27 and BICR56 cells, accumulation of acidic components and increased volume of acidic vesicular organelles was assessed by Acridin Orange (AO) staining. AO is a fluorophore that accumulates as protonated form in acidic vesicular organelles such as autolysosomes and is commonly used to study late-stage autophagy by fluorescence microscopy or flow cytometry [62]. As shown in **Figure S7A**, DX treatment alone caused only a small increase in AO fluorescence in SAS cells. DAS alone more than triples the percentage of autophagic cells. DAS+DX co-treatment further increased the proportion of cells with high fluorescence. Repeated measurements (n = 3) for all TSCC lines showed a similar response in all TSCC lines (**Figure S7B**). Upon DAS+DX co-treatment, an especially striking effect was seen for CAL27 cells, again pointing to pleiotropic effects of GCs when combined with DAS.

### The glucocorticoids Hydrocortisone and Methylprednisolone synergize with Dasatinib to suppress growth

Since many different glucocorticoids are in clinical use, we also wanted to explore whether other commonly prescribed drugs like Hydrocortisone (HC) and Methylprednisolone (MP) display DX-like activity in TSCC cells. For this, SAS cells were treated with DMSO (mock), 50 nM DAS, 300 nM HC or 150 nM MP, as well as combinations of DAS+HC or DAS+MP for 72 hours.

As **Figure 6A** shows, the cell density of SAS cells in 2D culture is clearly reduced after DAS treatment. This reduction is more pronounced after DAS+HC and DAS+MP combination treatments. To quantify cell numbers, cells were stained with trypan blue and counted. The cell number was reduced upon DAS treatment to ca. 65% of DMSO-treated cells. HC and MP treatment alone reduced viable SAS cells to approximately 80%. The antiproliferative activity of DAS was further increased after addition of HC or MP. The number of viable cells was reduced further from 65% to 48% following DAS+HC or to 42% following DAS+MP combination treatment (**Figure 6B**). In follow-up experiments, 3D soft agar colony formation was employed to examine the effects of HC or MP in a slightly more physiological context. Initially, SAS cells were treated with serial dilutions of HC or MP (**Figure 6C+D**). Data for CAL27 cells can be found in Supplementary **Figure S8A+B**. Colonies were counted after 14 days. As already seen for DX (**Figure 2C**), no noticeable decrease in colony formation was found for up to 10 µM with either of the GCs (**Figure 6C** for SAS and **Figure S8A** for CAL27).

**Figure 6.**
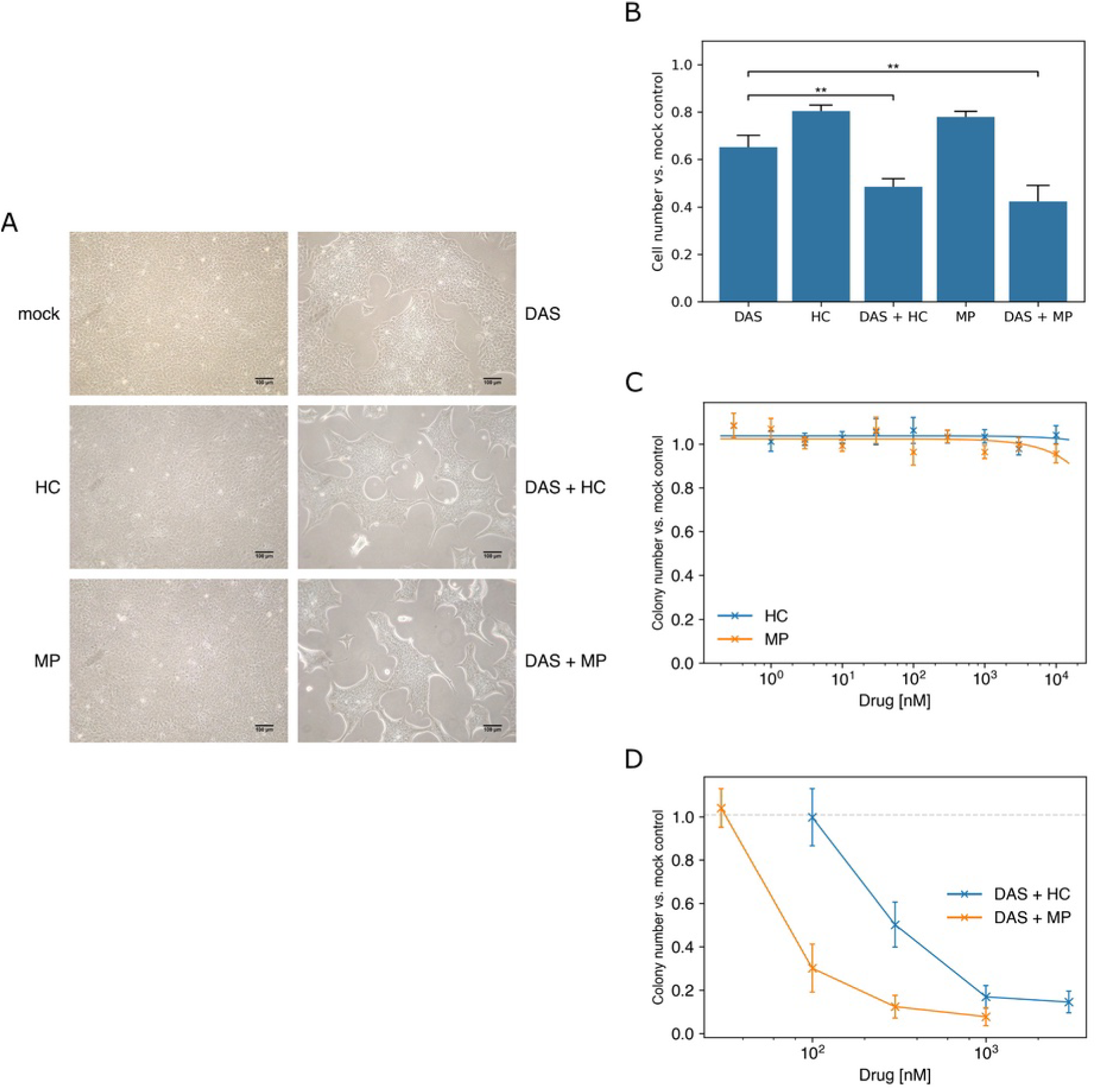
Hydrocortisone and Methylprednisolone show synergistic activity with Dasatinib. (A) SAS cells were treated with DMSO (mock), 50 nM of DAS, 300 nM of HC, 150 nM of MP, a combination of DAS+HC or a combination of DAS+MP for 72 h. Bright-field images of SAS cell growth in 2D monolayer culture after 72 h of treatment. Images of SAS cells were captured with an inverted microscope using 10x magnifications. (B) SAS cells were treated with DMSO (mock), 50 nM of DAS, 300 nM of HC, 150 nM of MP, a combination of DAS+HC or a combination of DAS+MP for 72 h. Viable SAS cell counts against mock control were evaluated by staining cells with trypan blue and counting of viable cells with an automated cell counter. Values are expressed as means and standard deviation obtained from three independent experiments. Asterisks indicate statistical significance for T-tests with p-values less than 0.01 (**). (C) HC and MP as single treatments show no reduction in the 3D soft agar colony formation of SAS cells as determined by colony counting. Data represent colony numbers normalized to mock-treated cells and show mean values ± SD of three independent experiments. (D) Concentration-dependent colony formation reduction of SAS cells in response to a fixed concentration of DAS (1 µM) and increasing concentrations of HC or MP in 3D soft agar culture. Data again represent colony numbers normalized to mock-treated cells and show mean values ± SD of three independent experiments.

Increasing concentrations of HC or MP were then combined with a marginally active concentration of 1 µM DAS for SAS cells (deduced from **Figure 2D**) or 100 nM for CAL27 cells (deduced from **Figure S4D**). DAS+HC and DAS+MP combination treatments exhibit strong colony formation reduction in a dose-dependent manner (**Figure 6D and S8B**). For HC, which is less potent than MP, a concentration of at least 300 nM was needed to show a notable effect in combination with DAS. MP displays a synergistic activity together with DAS at a concentration comparable to DX (100 nM). This indicates that at least four different GC show similar effects on TSCC growth in 2D and 3D culture. It should be noted that the investigated concentrations of HC and MP are pharmacologically achievable in human patient plasma upon normal dose administration [63, 64].

## Discussion and Conclusions

Molecularly targeted therapies are so far benefiting only a limited number of cancer patients. This is in part resulting from the high cost of these medicines [65], but also due to a limited arsenal of available drugs and our inability to reliably identify some of the patients who would significantly benefit from a particular type of targeted therapy [66]. The enormous heterogeneity of cancers on the molecular and biological level is at present only marginally understood. Even cancers arising in the same organ or body part can display an astounding degree of diversity. This has not yet been taken into sufficient consideration in the context of HNCs. Numerous preclinical and clinical studies have analyzed cancers originating from distinct sites of the head and neck as a more or less single entity rather than focusing on a specific tumor site [67]. However, epidemiological evidence and numerous studies by pathologists clearly show that major biological differences and clinical characteristics do exist between the different head and neck cancer sites and even sub-sites, as well as between different populations and individual patients with distinct underlying molecular defects [7, 68, 69]. Although there are now several targeted therapies for patients with HNSCC in clinical use [70], no particular targeted therapy for TSCC is available until now. More effective and more benign TSCC therapies are urgently needed. Identifying predictive tumor markers and new treatment targets, starting with cellular tumor models, is considered a valuable approach to achieving this.

Tyrosine kinases are frequent drivers of human tumor development but remain marginally studied in TSCC. Their mutation or hyper-activation often leads to constitutively elevated pTyr-protein levels that promote tumor progression. Our current preclinical approach, to find tyrosine kinases that are functionally implicated in the survival and growth of many different TSCC lines and to combine kinase inhibitor treatment by re-purposing other readily approved drugs, is hoped to be a rapid and cost-effective way forward in the quest for much better tumor treatments. Our starting point, a large panel of no less than 34 TSCC lines, was chosen in order to ensure that we focus on molecular targets shared by most, if not all, TSCC. Surprisingly, we found a high degree of similarity in the pTyr-protein patterns of our TSCC panel (**Figure 1A** and **Figure S1**). A broad 130 kDa pTyr-protein band immediately reminded us of p130Cas [23], the major substrate of the Src kinase, which was discovered as the first oncogene more than 100 years ago in the form of a tumor virus [71]. Strikingly, inhibition of SFKs had a massive impact on overall pTyr-protein levels as well as p130Cas phosphorylation, thereby documenting that SFKs are major drivers of tyrosine phosphorylation in TSCC lines. Alterations of p130Cas expression and interactions and the resulting regulatory signals are determinants for the occurrence of different types of human tumors [72, 73].

In the past years, several observations suggested that aberrant activation of the p130Cas signaling network signature leads to up-regulation of key regulatory signaling pathways promoting cell transformation. For example, p130Cas silencing in HER2-dependent breast carcinoma reduces HER2 levels and impairs migration and invasion *in vitro*, as well as the formation of lung metastases *in vivo* [74-76]. Taken together, these results suggested to us that DAS could be a useful drug for TSCC therapy.

However, since targeted therapies are very frequently of time-limited use due the development of tumor resistances, and DAS may not be sufficiently effective when applied as monotherapy in head and neck cancer patients [17], we decided to screen for FDA-approved drugs that can be used in combination with DAS. One of the substance classes that emerged from this screen and became the central focus of the current work are glucocorticoids. They are in abundant and manifold clinical use, inexpensive and well-characterized. Interestingly, the inhibitory effects seen with DAS+GC combinations tended to be more striking in the soft agar assay, which monitors anchorage-independent (3D) growth, than in regular 2D cell culture. This implies that we may not have seen some possibly useful drug combination synergies in our high-throughput screen, carried out in classical 2D culture.

One of the most striking results from our experiments is the profound, albeit probably indirect, effect of even low GC concentrations on MET - Gab1 signaling in TSCC lines. Clinical MET inhibition in various cancers is highly desirable and therefore an area of intense research, but so far hampered by significant toxicity and, at least in some cases, a lack of sufficient efficacy, of clinically tested MET-targeting kinase inhibitors [77-79]. More specifically, MET signaling is also known to play a role in HNCs with respect to tumor development, therapeutic resistance and the behavior of the tumor microenvironment [80]. All three analyzed TSCC lines display a basal activity of MET - Gab1 signaling (**Figure 4A**). This activity is sensitive even to subnanomolar concentrations of DX and FL (**Figure 4C**). The exact mechanism behind this remains unclear, as DX does not directly inhibit MET kinase activity (**Figure S5**).

Some reports of glucocorticoid activity on the HGF - c-Met - Gab1 signaling pathway have been previously published, including a link between GC and Gab1 signaling related to asthma [81]. GCs have also been shown to reduce HGF expression in cells for primary rat osteoblast cells [82]. This could potentially also occur in the TSCC lines, an avenue which remains to be explored.

Moreover, Gab1 has been shown to be important for the survival of HNSCC cells and a reduction of Gab1 expression led to impaired MAPK and PI3K - Akt signaling [83, 84]. This may, at least in part, be linked to the Gab1 effector phosphatase SHP2, since HNSCC cell lines were found to be particularly sensitive to the SHP2 inhibitor SHP099 when a large panel of over 800 cell lines of different tumor types was screened [84].

In any case, our current experiments have already provided several interesting leads into the pleiotropic functional biology of GCs in TSCC lines, which can now be followed up with more in-depth studies, in particular animal experiments. The full spectrum of bioactivity from DAS+GC treatments of TSCC lines will only become apparent once multiple signal transduction pathways and bioactivities are integrated and evaluated *in vivo*. The final functional outcomes of these integrative processes in the TSCC cell signaling networks are hard to predict. Consequently, the exact effects may vary from tumor to tumor, depending on the specific genetic and epigenetic alterations acquired during cancer development. Nevertheless, key results were found to be similar in three different cell lines, suggesting that they are likely to be representative for many other TSCC lines.

DAS, although clearly clinically useful and at present part of the standard anti-cancer drug portfolio in hospitals, may also be replaced in the future by more potent or more selective molecules, at least in some contexts. For example, the SFK inhibitor NXP900 [85], which is currently evaluated in early clinical trials, has shown striking activity in animal studies. NXP900 is reported to not only inhibit the kinase activity of SFK but also their kinase-independent scaffolding ability, increasing the potency of the inhibitor. Like DAS, its activity is enhanced when combined with other drugs. For example, NXP900 synergizes with the MEK inhibitor Trametinib in low grade serous ovarian cancer [86]. It is certainly worthwhile to test such newly arising, promising molecules in combination with different GCs for increased efficacy in TSCC in the near future. Moreover, the impact of GCs should be explored further in other squamous carcinomas of the head and neck region, in the hope of reducing the burden that our current HNC therapies elicit.

## List of abbreviations

AO: Acridin Orange
DAS: Dasatinib
CML: Chronic myelogenous leukemia
DX: Dexamethasone
EMT: Epithelial-mesenchymal transition
FL: Fluticasone
GC: Glucocorticoid
HC: Hydrocortisone
HNC: Head and neck cancer
HNSCC: Head and neck squamous cell carcinoma
HPV: Human papilloma virus
MAPK: Mitogen-activated protein kinase
MP: Methylprednisolone
OSCC: Oral squamous cell carcinoma
PTK: Protein tyrosine kinase
RTK: Receptor tyrosine kinase
SA-β-gal: Senescence-associated beta-galactosidase
SFK: Src family kinases
TSCC: Tongue squamous cell carcinoma

## Availability of data and materials

All data generated or analyzed during this study are included in this published article and its supplementary information files. The three cell lines mainly used (BICR56, CAL27, SAS) are available from commercial cell culture collections.

## Authors’ contributions

AH performed the major part of the biochemical experiments and analyses. JD investigated the patterns of phosphotyrosine proteins in the TSCC panel and performed experiments with SFK inhibitors. DE designed the HTP screen. SF conceived the project and supervised the work. ML performed the HTP screen, analyzed the data and supervised the work. AH, SF and ML drafted the manuscript. JD and DE commented on and approved the manuscript.

## Acknowledgements

We would like to thank Elena Seraia, formerly at the Target Discovery Institute, Oxford, UK, for supervising the practical part of the HTP screen and many interesting stories about life in Novosibirsk. We would also like to thank Nadine Bley, Core Facility Imaging (Martin-Luther-University Halle-Wittenberg, Germany), for her help in performing the flow cytometry experiments and the data analysis.

## Methods

### Reagents

PP2 (#529573) and PP3 (#529574) were purchased from Calbiochem, Dasatinib (#HY-10181) from Hycultec, and Crizotinib (#S1068) from Selleckchem. Dexamethasone (#D4902), Hydrocortisone (#H4001) and Methylprednisolone (#S1733) were obtained from Sigma-Aldrich. Antibodies against Src (#2108), pTyr416 Src (#2101), pTyr410 p130Cas (#4011), CDK1 (#9116), CDK2 (#2546), CDK4 (#12790), CDK6 (#13331), Cyclin B1 (#12231), WEE1 (#4936), pSer462 WEE1 (#4910), MET (#3148), pTyr1234/1235 MET (#3077), pTyr627 Gab1 (#3233) and pThr202/Tyr204 ERK 1/2 (#9101) were bought from Cell Signaling Technologies. Antibodies against p130CAS (#610271), Cyclin D3 (#610279) and p27Kip1 (#610241) were acquired from BD Biosciences. The antibody against Actin was purchased from Sigma-Aldrich (#A5441), anti-ERK 1/2 (#06-182) from Upstate Biotechnology and anti-Gab1 (#A303-288A) from Bethyl Laboratories. Secondary antibodies, HRP anti-mouse IgG (#715-036-151) and HRP anti-rabbit IgG (#711-036-152), were bought from Jackson ImmunoResearch Labs.

### Cell lines and culture conditions

The origin of the 34 TSCC lines used in this study is described in **Supplemental Table S1** and the literature [22]. All cell lines were initially established from primary tumors, except for the HSC-3, HSC-4, OSC-19 and OSC-20 cell lines, which were established from metastatic sites. Cells were cultured in Dulbecco’s Modified Eagle Medium with high glucose and glutamine (DMEM, Gibco, #41966-029) supplemented with 10% heat-inactivated fetal bovine serum (FBS, Biowest, #S1860-500), 1% non-essential amino acids (NEAA, Gibco, #11140-035), and 1% penicillin/streptomycin (Gibco, #15140-122) at 37 °C in a humidified atmosphere with 10% CO_2_. Cells were grown to 85-90% confluency and then passaged by trypsinization to avoid overgrowth.

### Western blot analyses

TSCC cells were plated in 10-cm culture dishes and treated with the indicated compounds and concentrations. At the end of the treatments, proteins were extracted with RIPA lysis buffer (20 mM Tris-HCl pH 7.5, 1 mM EDTA, 100 mM NaCl, 1% (v/v) Triton X-100, 0.5% (w/v) DOC and 0.1% (w/v) SDS) in the presence of protease (1:40 cOmplete™ Protease Inhibitor Cocktail (Roche, #11836145001)) and phosphatase inhibitors (1:100 Phosphatase inhibitor cocktail 2 (Sigma Aldrich, #P5726), 1:100 Phosphatase inhibitor cocktail 3 (Sigma Aldrich, #P0044)). Protein concentrations were determined according to the method of Bradford. Equal amounts of proteins were separated by SDS-PAGE and transferred to a PVDF membrane by semi-dry transfer blotting. The membrane was then incubated for 1 h at room temperature (RT) in a blocking buffer (5% of BSA or milk in TBST (20 mM TrisHCl pH 7.5 and 100 mM NaCl with 0.1% (v/v) Tween 20)) as appropriate for the individual antibody. It was then incubated with the respective primary antibody in blocking buffer overnight at 4 °C on a nutator. After washing three times with TBST, the membrane was incubated with horseradish peroxidase (HRP)-coupled secondary antibody for 1 h at RT. After washing three more times with TBST, a freshly mixed enhanced chemiluminescence (ECL) detection solution (Advansta, #K-12043-D10) was applied. The protein signals were detected on a chemiluminescence imaging system (Syngene G:BOX, Avantor).

### MET kinase assays

SAS cells were seeded in 10 cm cell culture dishes, grown for 24 h and lysed with kinase lysis buffer (50 mM TrisHCl pH 7.5, 150 mM NaCl, 1% (v/v) Triton X-100, 1 mM EGTA, 1 mM EDTA and 1% (v/v) Glycerol) containing the appropriate inhibitors (0.5 ng/ml Leupeptin, 10 ng/ml Aprotinin, 200 ng/ml PMSF (DMSO), 1 mM Sodium orthovanadate, 1:100 Phosphatase inhibitor cocktail 2, 1:100 Phosphatase inhibitor cocktail 3 and 1:40 cOmplete™ Protease Inhibitor Cocktail). The cell lysate was agitated at 4 °C for 1 h, centrifuged and the supernatant was transferred to a new tube. The protein concentration was determined by Bradford assay. For pre-clearing, 10 µl amounts of G Sepharose beads (Cytiva, #17061801) were transferred into 1.5 ml tubes, washed twice with 500 µl of lysis buffer and mixed with 500 µg of protein cell lysate on a nutator at 4 °C for 1 h. Samples were centrifuged, the beads were discarded and the pre-cleared supernatants were transferred into new tubes. Subsequently, three types of immunoprecipitations were prepared (Table 1). 5 µg of mouse isotype control (Cell Signaling Technology, #5415) or 5 µg of MET antibody (Cell Signaling Technology, #8198) were added into the tubes containing pre-cleared lysates. Samples were agitated for 2 h at 4 °C, 10 µl G Sepharose beads were added and then mixed for 1 h at 4 °C. Then, the samples were centrifuged and the supernatants were discarded. The immunocomplexes were 3x washed with 500 µl of washing buffer (20 mM TrisHCl pH 7.5, 100 mM NaCl, 1% (v/v) Triton X-100, 1 mM EGTA, 1 mM EDTA and 2.5% (v/v) glycerol) and kept on ice.

**Table 1.** Types of samples generated for immunoprecipitation (IP)

| Sample | Isotype control | No lysate control | Immunoprecipitation (IP) |
| --- | --- | --- | --- |
| Lysis buffer | X | same volume as lysate | X |
| Lysate | 500 µg | X | 500 µg |
| Antibody | 5 µg isotype control | 5 µg MET antibody | 5 µg MET antibody |

Samples were prepared as described in Table 2, added onto the immunocomplexes and incubated in a thermomixer (22 °C, 500 rpm) for 5 min. 5 µl of 20 mM ATP (pH 7.0) was added to all of the samples (except for the ‘No ATP control’) to start the kinase reaction. Samples were again incubated in a thermomixer (22 °C, 500 rpm) for 30 min and the kinase reaction was stopped by addition of 50 µl SDS-PAGE sample buffers. Samples were immediately boiled at 95 °C for 15 minutes and centrifuged. 7.5 µg of the total cell lysate and 30 µl of the supernatants were separated by SDS-PAGE and analyzed by Western blotting as described above with pTyr627 Gab1, total MET and pTyr1234/1235 MET antibodies.

**Table 2.**
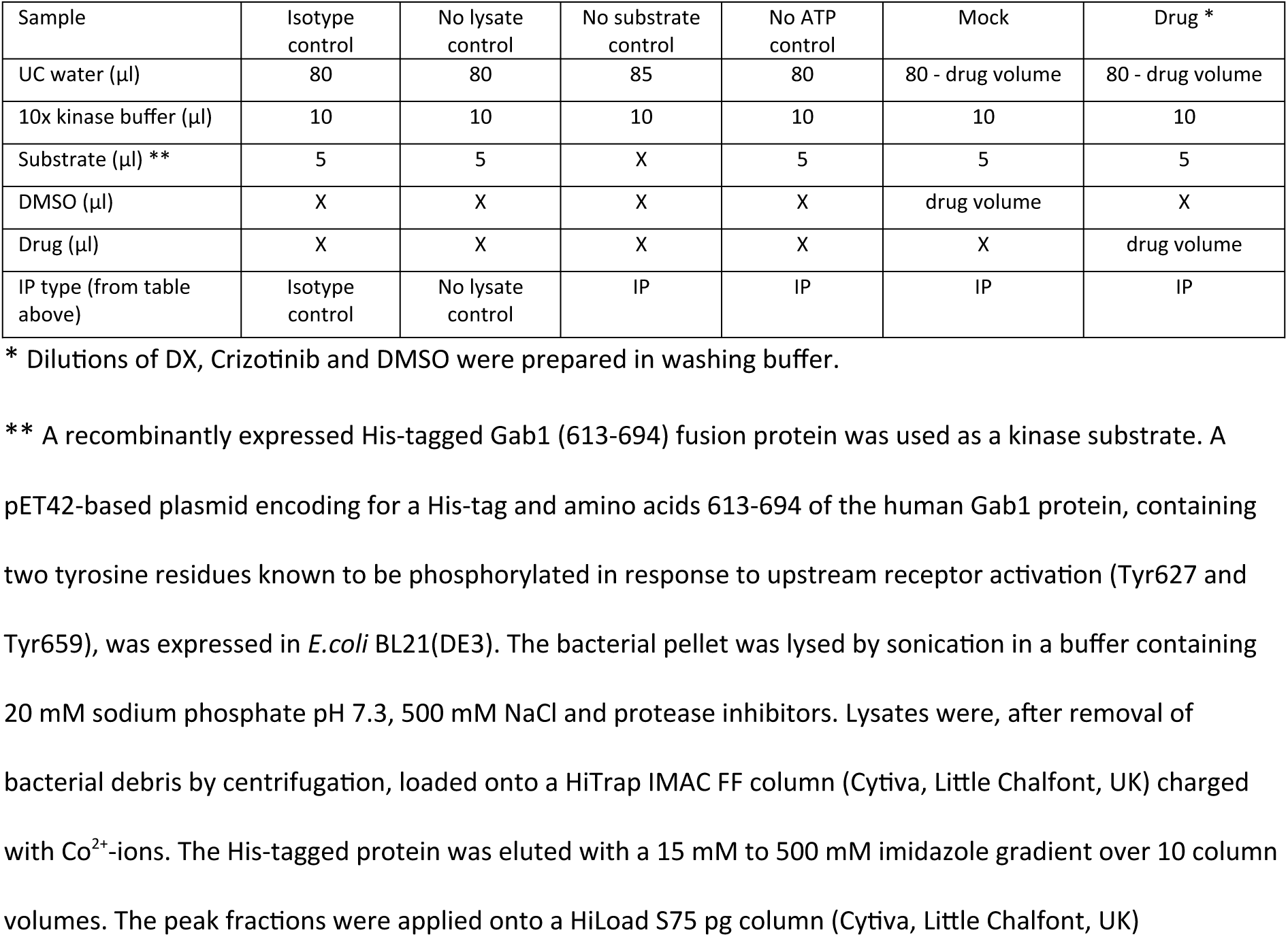
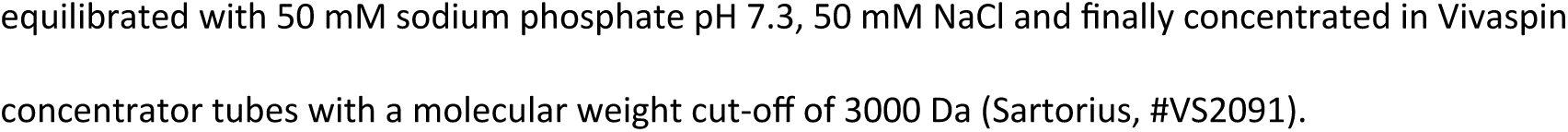
Sample preparation for the MET kinase assay.

### Resazurin cell viability assays

The cytotoxicity of all compounds in 2D culture was evaluated by a resazurin assay. TSCC cells were washed with PBS, trypsinized, counted and seeded into 96-well plates at the appropriate cell densities to prevent confluence of the cells during the period of the experiment. For SAS, CAL27, and BICR56 cell lines 8000, 5000, and 3500 cells/well were seeded, respectively. After 24 h, the cells were treated with the investigated compounds or DMSO as vehicle control (mock). After treatment, the medium was discarded and each well was washed with 100 µl of PBS. The wells were then stained with 100 μl of 0.1% (w/v) resazurin (Sigma Aldrich, #199303) in DMEM without phenol red (Gibco, #21063-030) for about 2 h at 37 °C. The fluorescence at 590 nm was measured after excitation at 531 nm using a Infinite M Plex plate reader (Tecan, Männedorf, Switzerland). The percentages of surviving cells related to mock-treated cells were determined. Each experiment was repeated at least 3 times. Results are presented with the mean ± standard deviation.

### Trypan Blue exclusion assays

The Trypan Blue dye exclusion assay was used to determine the number of viable and dead cells in a cell suspension. For this, cells were plated in 6-well plates at the suitable cell numbers per well. For SAS, CAL27, and BICR56 cell lines, 240,000, 150,000, and 100,000 cells/well were seeded, respectively. On the following day, cells were exposed to different concentrations of agents or DMSO as vehicle control (mock). After 72 h of drug exposure, 10 µl of cell suspension was mixed with 10 µl of Trypan Blue (Invitrogen, #T10282) and pipetted into a disposable Countess cell counter chamber slide (Invitrogen, Waltham, MA, USA). Cell numbers were determined with an automated Countess II FL cell counter (Invitrogen, Waltham, MA, USA). Cell viability was calculated using the ratio of total live cells normalized to total live cells in the mock-treated sample. Each experiment was replicated three times.

### Morphological evaluation by phase-contrast inverted microscope

Microscopic investigation was performed to identify the morphological features of treated cancer cells in comparison to mock-treated controls. For this, cells were seeded in 6-well plates and then treated with different concentrations of the investigated compounds or DMSO as vehicle control (mock). After 72 h of drug exposure, cell growth was evaluated with an inverted microscope (Eclipse TS100, Nikon) and photos were captured with a digital camera (Coolpix E5400, Nikon) at 10x and 20x magnifications.

### 3D soft agar assays

The anchorage-independent colony formation assay (soft agar assay) was performed as described previously [87]. 0.6% (w/v) Noble agar (BD Biosciences, #214220) in 1x DMEM was plated as a bottom layer into 48-well plates placed on a pre-warmed glass plate (42 °C). Only the inner 24 wells of the 48-well plates were used, to avoid edge effect. Plates were left to harden for at least 30 min at RT in the cell culture hood. During that time, cells were trypsinized, diluted with 1x DMEM and counted. The cells were diluted with DMEM and mixed into 0.6% agar for a final agar concentration of 0.4% (6,000 cells per well). This second layer was plated on top of the bottom (0.6% agar) layer. The plates were left again for at least 30 min at RT to solidify.

After solidification, 200 µl of DMEM with DMSO or compounds was added into each well. The cells were cultured at 37 °C in 10% CO_2_ and the wells were fed every 3 to 4 days with 70 µl of DMEM with DMSO or compounds, stored since day 1, to prevent desiccation. After 14 days of incubation, cells were stained overnight at 37 °C by adding 70 μl of 0.5% nitroblue tetrazolium chloride (Roth, #4421.3) solution per well. Once colonies were stained, photographs of wells were taken using a Sony Alpha 5100 camera with a SEL30M35 macro lens. The colonies were counted using image analysis software (ImageJ 1.50i) using a minimum diameter of 50 µm as a cut-off for viable colonies. All experiments were conducted in triplicates. Calculation of IC_50_ values was performed in R 4.2.3. Values are presented as the means ± SEM from three independent experiments.

### Cell cycle analyses

TSCC cells were seeded in 6-well plates for 24 h and treated with the indicated concentrations of each compound or DMSO as vehicle control (mock) for 72 h. The supernatant with the floating dead cells and the attached cells from each well were collected and fixed in 3 ml of cold ethanol at -20 °C overnight. Cells were stained with DAPI (1:10,000 dilution of a 20 mg/ml DAPI (Sigma Aldrich, #D9542) stock and 1:5,000 of 10 mg/ml RNAse A in PBS) for 30 min at 37 °C in the dark and then washed with PBS. Cell fluorescence was measured with a MACSQuant Analyzer 10 (Miltenyi Biotec) flow cytometer. Histograms of cell numbers vs. blue fluorescence were recorded and the distribution of cells in different cell cycle phases was determined based on the mean fluorescence intensity values. A total of 10,000 events were analyzed using the FlowJo (version 10) software.

### Analysis of autophagy by Acridine Orange staining

Autophagy was investigated by the staining of acidic vesicular organelles with Acridine Orange (AO). TSCC cells were trypsinized, counted, seeded in 6-well plates for 24 h and then treated with the indicated concentration of each compound or DMSO as vehicle control (mock) for 72 h. Cells were then collected, washed with PBS and stained with 500 µl of 10 μM AO solution (Sigma Aldrich, #235474) for 15 min at 37 °C in the dark. Afterwards, cells were washed with PBS to remove excess AO and the cell pellet was resuspended in 500 μl PBS for flow cytometry analysis with the MACSQuant Analyzer 10 (Miltenyi Biotec) flow cytometer. A total of 10,000 events were analyzed using the FlowJo (version 10) software.

### Senescence detection by β-Galactosidase staining

To detect the senescence state of the cells, treated TSCC were stained using a Senescence associated β-galactosidase (SA-β-gal) Staining Kit (Cell Signaling Technology, #9860) according to the manufacturer’s instructions.

Briefly, TSCC cells were seeded in 6-well plates for 24 h and treated for 72 h with the investigated compounds or DMSO as vehicle control (mock). The growth medium was discarded, the cells were washed with PBS and fixed in 2% paraformaldehyde for 15 minutes at RT. After washing twice with PBS, they were incubated overnight with the β-Galactosidase staining solution at 37 °C in a dry incubator without additional CO_2_. On the next day, while the β-Galactosidase was still on the plate, the cells were analyzed under the Eclipse microscope for the development of blue color and photographed with a digital camera (Coolpix E5400, Nikon).

### Life cell imaging

SAS cells were seeded at 150,000 cells per well into 24-well plates and left to attach for 24 h. The medium was then replaced with fresh medium containing 0.05% DMSO or 30 µM PP2. The plates were transferred into the Cell-IQ imaging system (Chip-Man Technologies Ltd). Photographs were taken at the same location every 20 min over 5 days. The Cell-IQ ImgenTM Software Package and Cell-IQ Analyser® software package were used to analyze the images and to create movie files.

### High-throughput screening and analysis

Three cell lines (BCR56, CAL27, and SAS) were used for high-throughput screening. Their identity was confirmed by STR profiling (Microsynth AG, Balgach, Switzerland) and the cells were free of mycoplasma as determined by PCR testing [88]. The cells were treated with a combination of DAS (0 µM or 0.03 µM) and FDA-approved drugs from the Pharmakon 1600 library (MicroSource Discovery Systems, Inc.; Gaylordsville, CT, USA) at three concentrations (0.4 µM, 2 µM, 10 µM). There was one biological replicate per treatment.

The compound library consisted of 20 library plates with 80 compounds each. The treatment was performed on seven different days as a maximum of three library plates could be processed within one day.

In preparation for each treatment day, cells were thawed from a frozen stock and passaged twice. The cells were then trypsinized, seeded into 96-well plates at 2700 (SAS), 4700 (CAL27) or 3400 (BICR56) cells/ well using a Flexdrop Plus dispenser (PerkinElmer, Waltham (MA), USA) and grown for one more day at 37 °C and 10% CO_2_. Compounds and controls were applied onto 96-well plates in a Janus MDT workstation (PerkinElmer, Waltham (MA), USA). On all plates, four wells each were reserved for medium-only, Staurosporine, DMSO, and DAS-only controls. All plates were prepared in duplicate.

The medium was replaced after three days on the Janus MDT workstation against DMEM without phenol red but containing resazurin (100 µg/ml). After another 2 h incubation, the metabolic activity was assessed by fluorescence measurement (excitation 531 nm, emission 595 nm) on an Envision Multilabel Reader (PerkinElmer, Waltham, MA, USA). r^2^ values (linear regression) were determined for plate duplicates. Plates with an r^2^ of less than 0.7 were excluded from further analysis (26 of 720 plates). Ratios for the two replicate wells, on different plates, were calculated. Outliers (1.5x interquartile range) were excluded from further analysis (45 of 34560 pairs). All sample wells on each plate were then normalized with the mean of their four DMSO-control wells. The mean of the two replicate sample wells from these DMSO-normalized plates was used for further analysis.

Ratios of cells treated with a combination of compound and DAS against the compound alone were log_2_-transformed for an approximately normal distribution and analyzed for outliers (1.5x interquartile range). Drugs that showed potential synergy in more than one cell line were considered to be the most promising candidates for further analyses.

### Statistical analyses

Data are presented as mean ± standard deviation, unless indicated otherwise. Model fits for EC_50_ estimation and standard error calculation were performed in R 4.2.3 [89] using the ‘drc’ 3.0-1 package ‘LL.4’ four-parameter log-logistic function [90]. T-tests were calculated with pingouin 0.5.4 [91]. Further statistical analysis was done with pandas 2.0.3 [92] and numpy 1.24.3 [93]. Charts were generated in Python 3.11.5 using the matplotlib 3.7.2 [94] and Seaborn 0.12.2 [95] libraries.

## Supplementary figure legends

**Figure S1.**
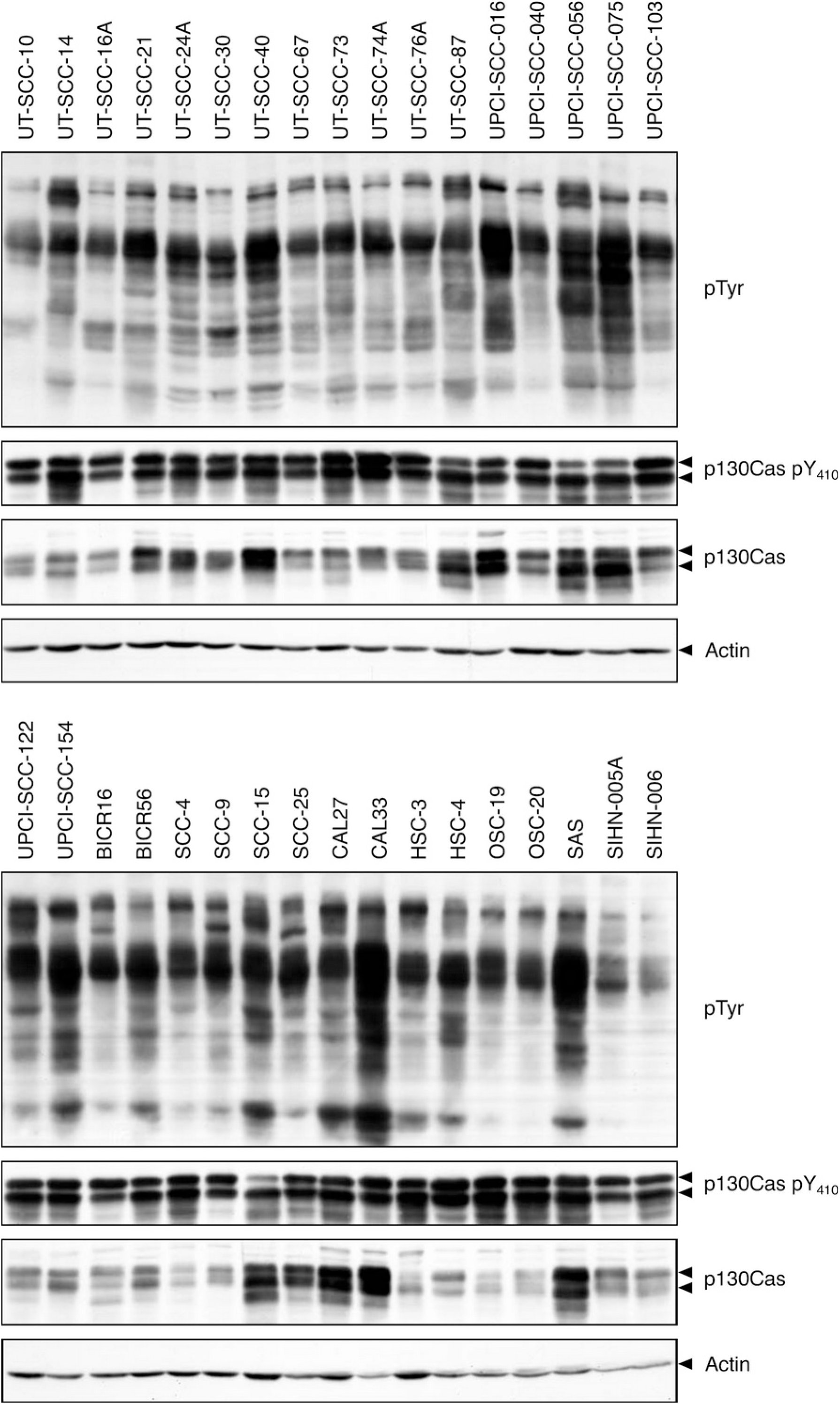
pTyr-profiling of 34 TSCC lines shows a uniform pTyr pattern and a prominent 130kDa band. Total cell lysates were analyzed by Western blotting for abundance and size of pTyr proteins using an anti-phosphotyrosine antibody (4G10). The expression levels of p130Cas and the phospho-status of p130Cas Tyr410 were also determined. Actin blots serve as loading controls.

**Figure S2.**
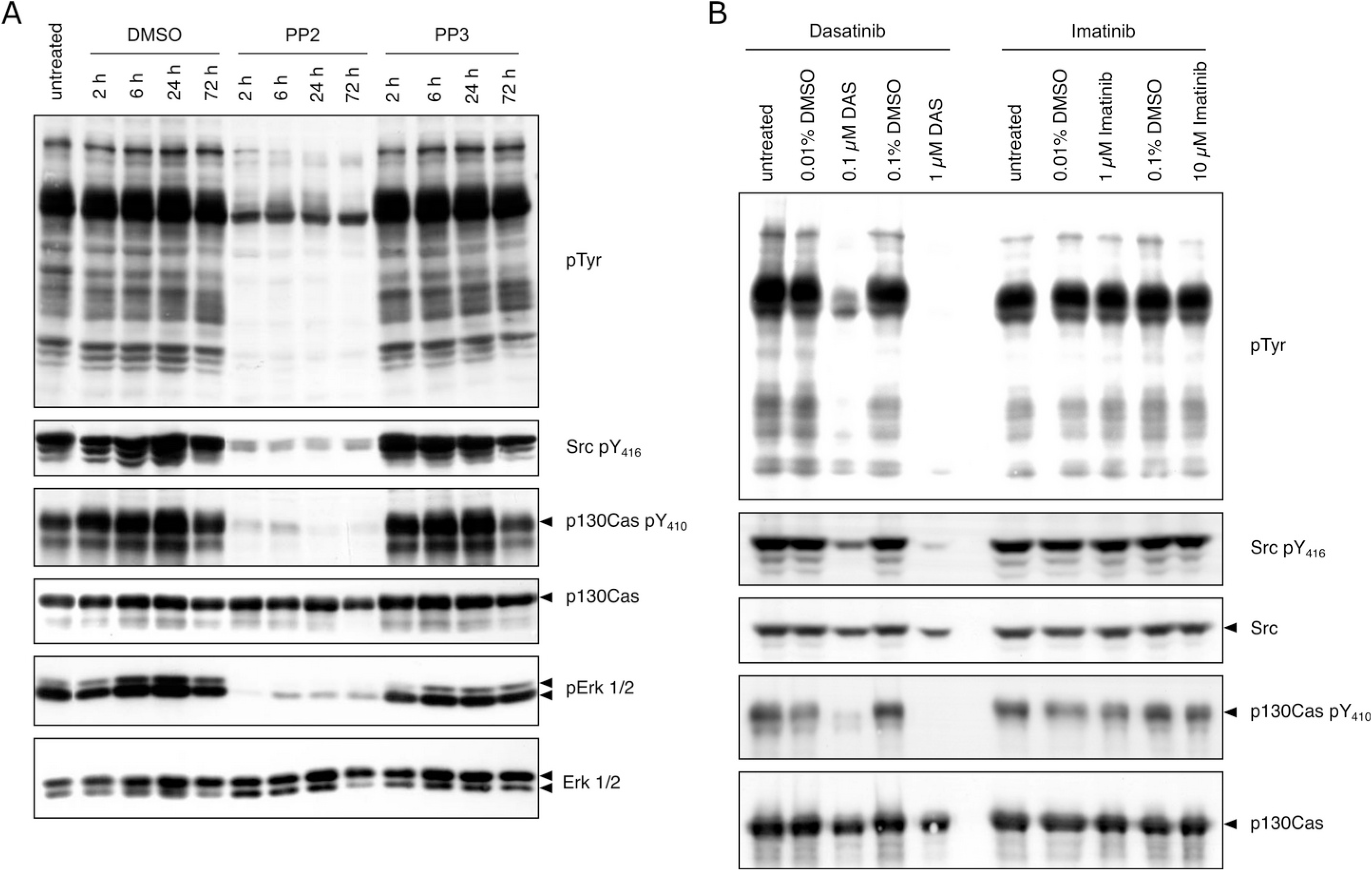
Src family kinase but not Abl inhibition reduces pTyr levels, p130Cas phosphorylation and activities of SFK and Erk1/2. (A) SAS cells were treated for up to 72 h, as indicated, with DMSO as vehicle control, 30 µM of the SFK inhibitor PP2 or 30 µM of the structurally similar control compound PP3, which lacks activity against SFK. Cells were then harvested and lysates were analyzed by Western blotting for pTyr patterns, p130Cas expression, phosphorylation on p130Cas Tyr410, as well as protein levels and activation of Erk1 and 2 as indicated. (B) SAS cells were treated with clinically relevant concentrations of the pleiotropic kinase inhibitors DAS (0.1 µM or 1 µM) or Imatinib (1 µM or 10 µM) or with DMSO as indicated. After 24 h, cells were lysed in SDS-PAGE sample buffer and analyzed by western blot for pTyr protein levels, p130Cas phosphorylation, SFK activity (pSFK) and Src expression. Phosphorylation of c-Src (chicken) on Y416 (pY419 in human) in the kinase activation loop is essential for Src activity, the epitope is conserved throughout evolution and different members of the SFKs [23].

**Figure S3.**
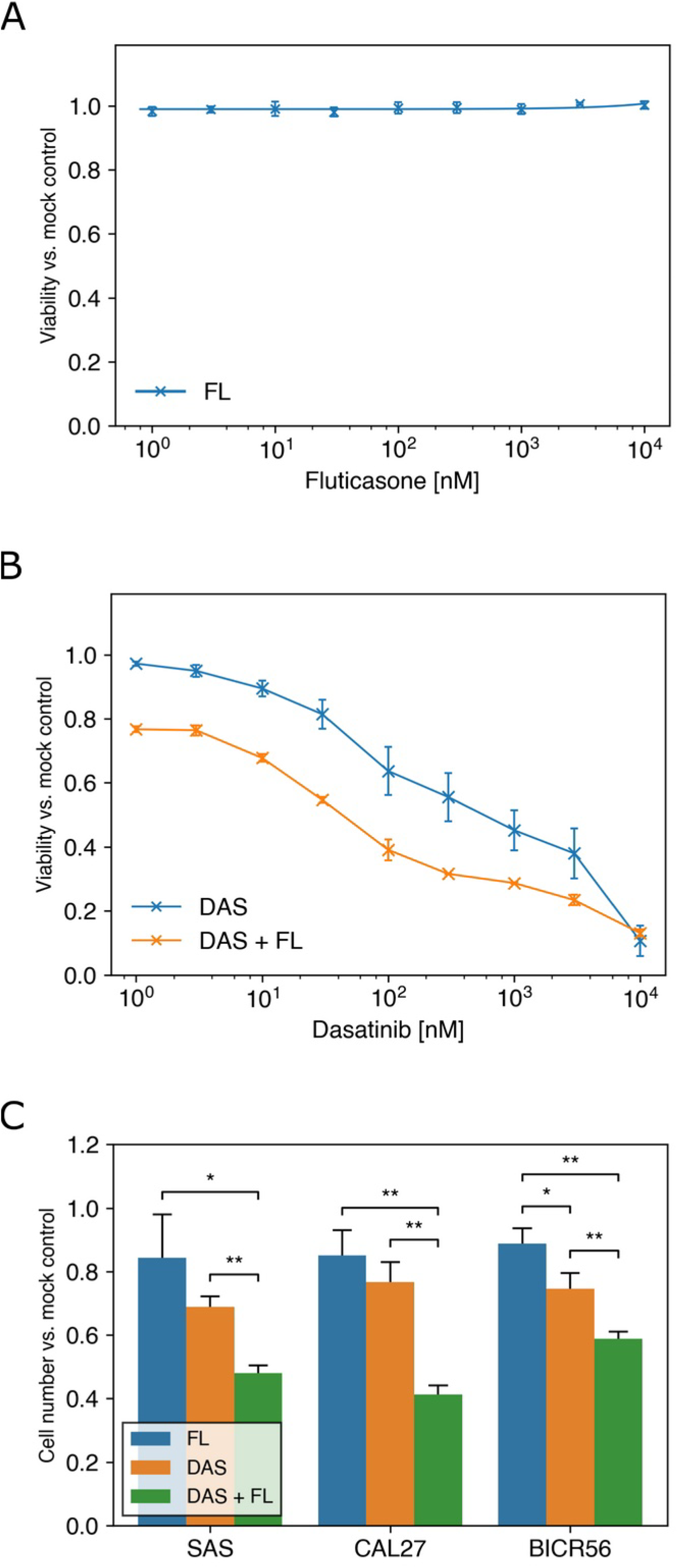
Addition of Fluticasone to Dasatinib-treated cells reduces cell viability and proliferation. (A) At a concentration of 10 µM, FL alone shows no notable effect on SAS cell line viability in 2D culture after 72 h treatment as measured via resazurin assay. (B) SAS cells were treated with increasing concentrations of DAS with or without 25 nM of FL for 72 h. Cell viability after treatment was measured by resazurin assay. (A and B) Data represent fluorescence measurements normalized to mock-treated cells. The dose-response curves show mean values ± SD of three independent experiments with six parallel measurements each. (C) TSCC cells were treated with DMSO (mock), 50 nM of DAS, 25 nM of FL, or a combination of DAS+FL for 72 h. Cell proliferation was assessed by staining cells with trypan blue and counting viable cells by automated cell counter. Cell numbers were normalized to mock-treated cells. Values are expressed as means and standard deviations obtained from three independent experiments. Asterisks indicate statistical significance for T-tests with p-values at less than 0.05 (*) or less than 0.01 (**), respectively.

**Figure S4.**
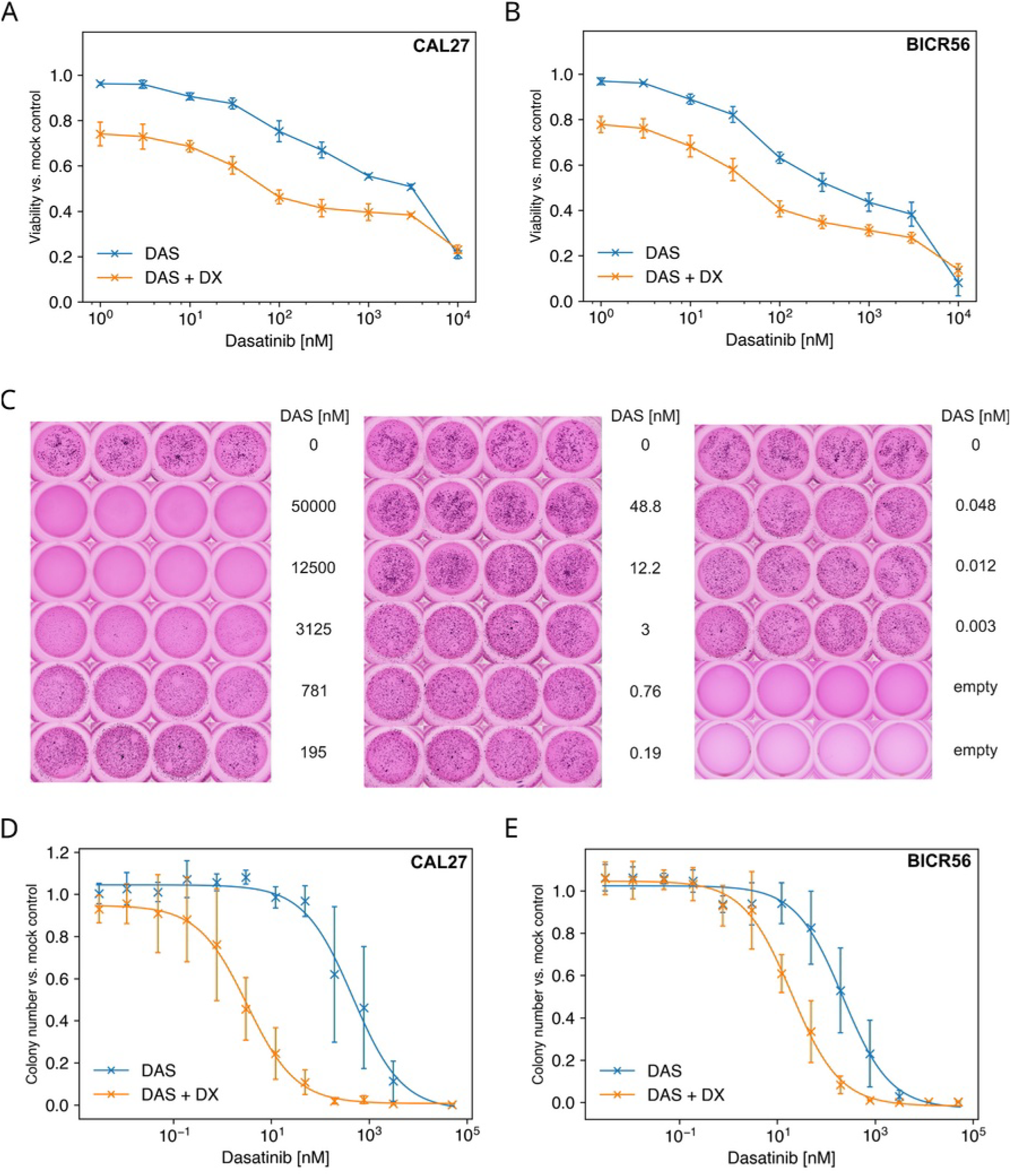
Synergistic activity on CAL27 and BICR56 is seen after DAS+DX combination in 2D and 3D. (A and B) Dose-response curves of CAL27 cells and BICR56 cells in 2D culture treated with different concentrations of DAS with or without 100 nM of DX for 72 h. Cell viability after treatment was measured by resazurin assay. (C) Representative results for a 3D soft agar experiment. SAS cells were treated with DAS concentrations from 0.003 nM to 50 µM. Colonies were stained after 14 days of incubation with nitroblue tetrazolium chloride and photographed. (D and E): Dose-response curves of CAL27 cells and BICR56 cells treated with DAS with or without DX (100 nM), as determined by colony numbers in 3D soft agar culture. In A, B, D and E, values are normalized to mock-treated cells and the dose-response curves show mean values ± SD of three independent experiments.

**Figure S5.**
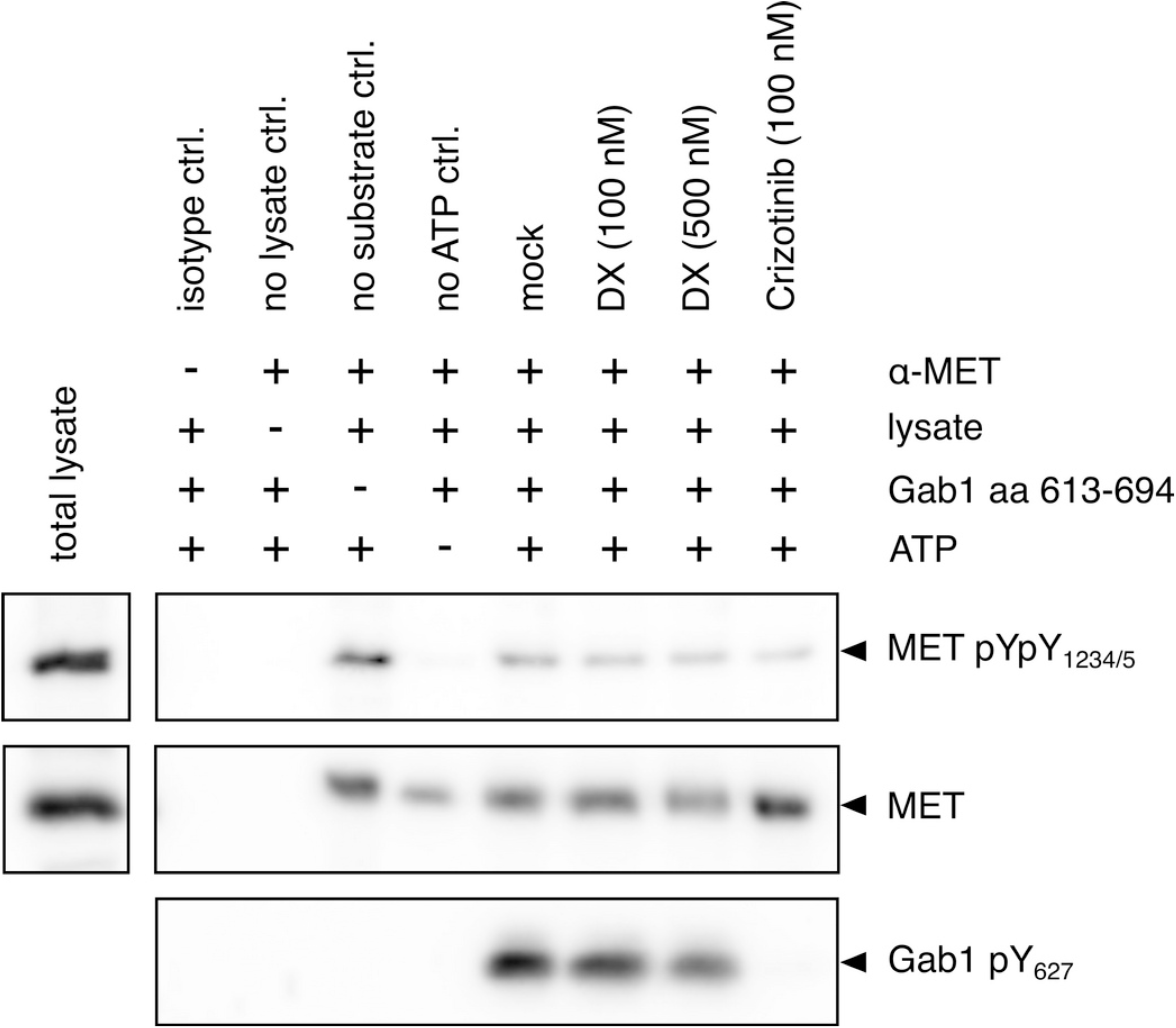
Dexamethasone shows no direct inhibition of MET kinase activity. MET kinase was immunoprecipitated from untreated SAS cells. The immunocomplexes were used in a kinase assay with a recombinantly expressed fragment of Gab1 (aa 613-694) containing two putative tyrosine phosphorylation sites (Tyr627 and Tyr659) as a kinase substrate. DMSO, 100 nM of DX, 500 nM of DX or 100 nM of Crizotinib (positive MET inhibitor control) were added to the samples. As further controls an antibody isotype control, a ‘no lysate’ control, a ‘no substrate’ control and a ‘no ATP’ control were included. Lysates were separated by SDS-PAGE and immunoblots were probed with the indicated antibodies.

**Figure S6.**
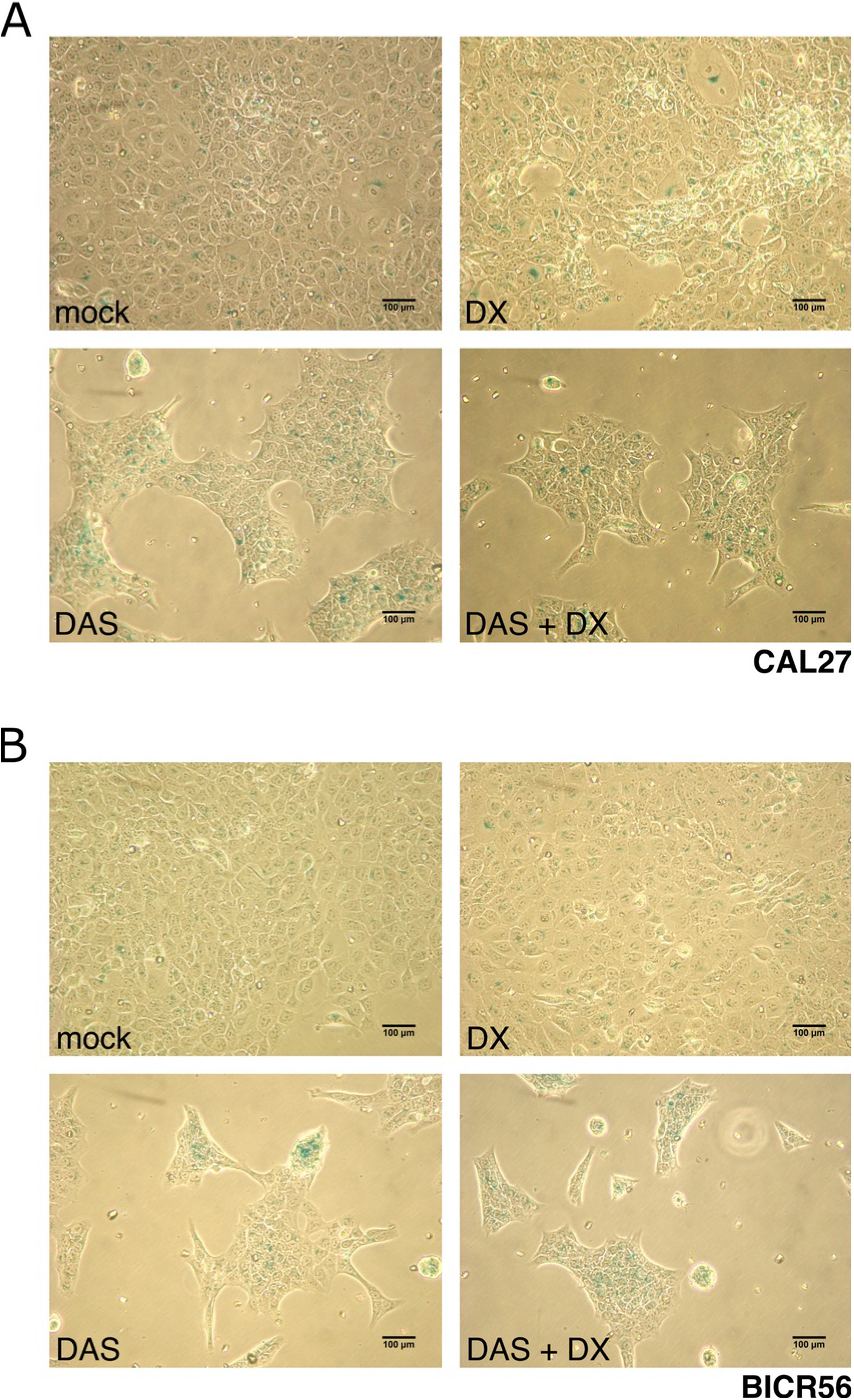
Dasatinib + Dexamethasone combination treatment appears to increase senescence in CAL27 and BICR56 cells. TSCC cells were treated with DMSO (mock), 50 nM of DAS, 100 nM of DX or a combination of DAS+DX for 72 h and stained with senescence-associated-β Galactosidase (SA-β-Gal). SA- β-Gal positive staining (blue color) indicates aged cells. Results for CAL27 (A) and BICR56 (B) are shown. The scale bar indicates a length of 100 µm.

**Figure S7.**
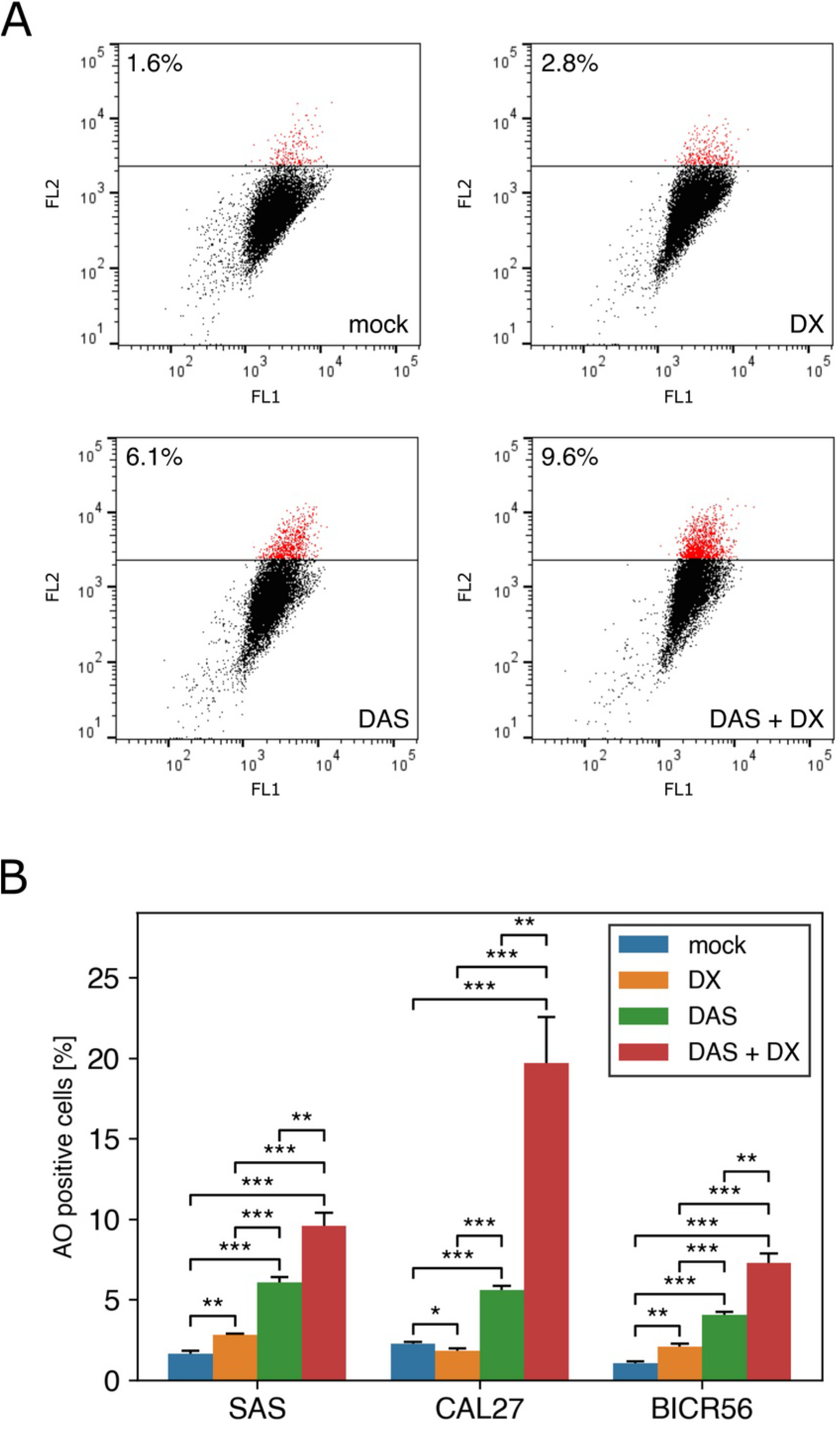
Dasatinib + Dexamethasone combination induces autophagy in TSCC lines. (A and B): TSCC cells were treated with DMSO (mock), 50 nM of DAS, 100 nM of DX, or a combination of DAS+DX for 72 h, stained with acridine orange (AO) and analyzed by flow cytometry to quantify the acidic vesicular organelle formation. (A) FL1 (x-axis) indicates green color intensity (530 nm wavelength), while FL2 (y-axis) shows red color intensity (680 nm wavelength). Representative results from three independent experiments are shown. (B) The AO-positive cells were quantified and shown as a percentage of the total cell population. Values are expressed as means and standard deviation obtained from three independent experiments.

**Figure S8.**
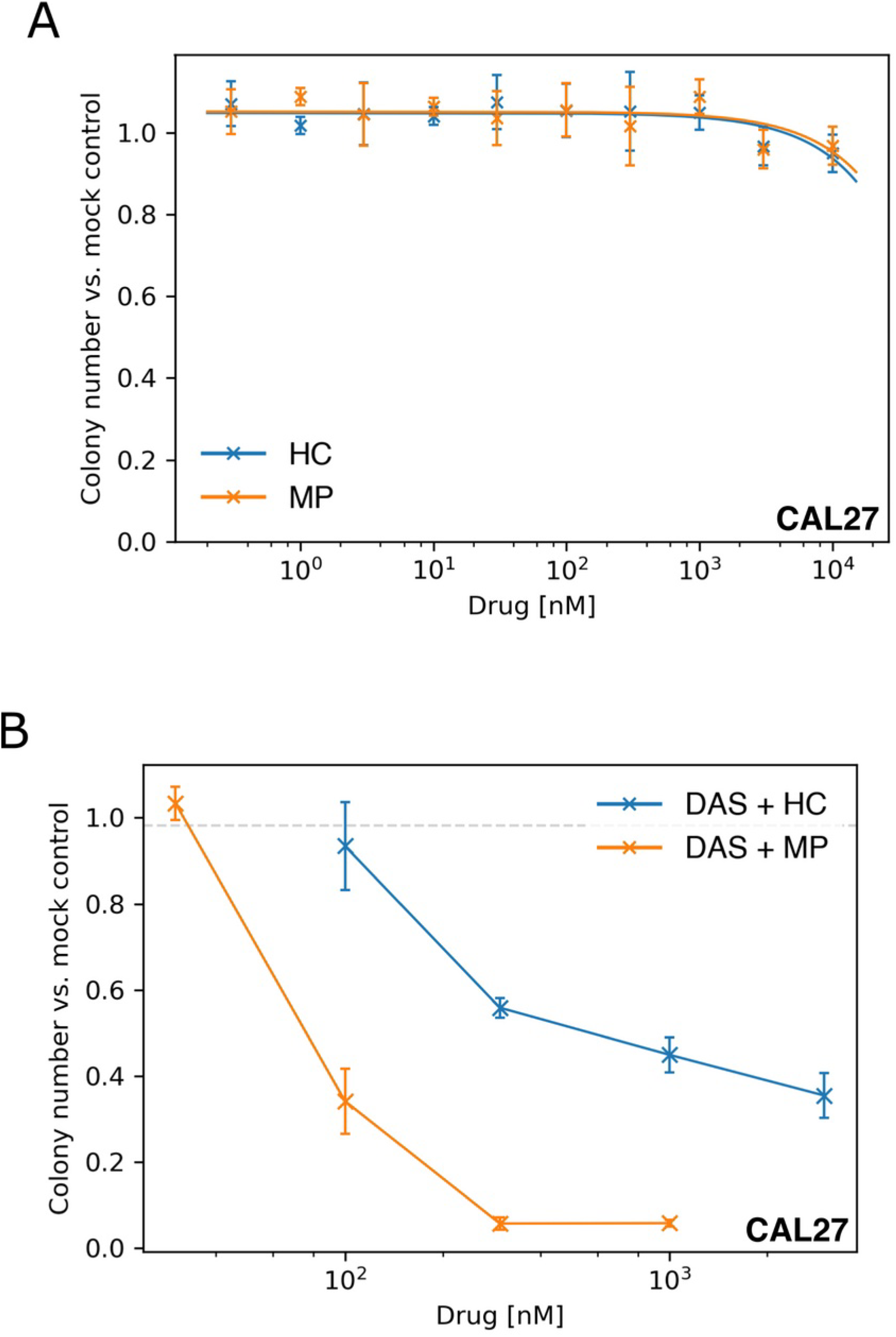
Hydrocortisone and Methylprednisolone show promising synergistic activity with DAS in CAL27 cells. (A) Dose-response curves of CAL27 cells treated with different concentrations of HC or MP as assessed by 3D soft agar colony formation. (B) Colony formation reduction of CAL27 cells in response to a fixed concentration of DAS (100 nM) and increasing concentrations of HC or MP in 3D soft agar culture. (A and B) data represent colony numbers normalized to mock-treated cells and the results show mean values ± SD of three independent experiments.

## References

1. Tan Y, Wang Z, Xu M, Li B, Huang Z, Qin S, Nice EC, Tang J, Huang C: Oral squamous cell carcinomas: state of the field and emerging directions. Int J Oral Sci 2023, 15:44.

2. Harada H, Kikuchi M, Asato R, Hamaguchi K, Tamaki H, Mizuta M, Hori R, Kojima T, Honda K, Tsujimura T, et al: Characteristics of oral squamous cell carcinoma focusing on cases unaffected by smoking and drinking: A multicenter retrospective study. Head Neck 2023, 45:1812–1822.

3. Campbell BR, Netterville JL, Sinard RJ, Mannion K, Rohde SL, Langerman A, Kim YJ, Lewis JS, Jr., Lang Kuhs KA: Early onset oral tongue cancer in the United States: A literature review. Oral Oncol 2018, 87:1–7.

4. Ramqvist T, Grün N, Dalianis T: Human papillomavirus and tonsillar and base of tongue cancer. Viruses 2015, 7:1332–1343.

5. Satgunaseelan L, Allanson BM, Asher R, Reddy R, Low HTH, Veness M, Gopal Iyer N, Smee RI, Palme CE, Gupta R, Clark JR: The incidence of squamous cell carcinoma of the oral tongue is rising in young non-smoking women: An international multi-institutional analysis. Oral Oncol 2020, 110:104875.

6. Vettore AL, Ramnarayanan K, Poore G, Lim K, Ong CK, Huang KK, Leong HS, Chong FT, Lim TK, Lim WK, et al: Mutational landscapes of tongue carcinoma reveal recurrent mutations in genes of therapeutic and prognostic relevance. Genome Med 2015, 7:98.

7. Leemans CR, Snijders PJF, Brakenhoff RH: The molecular landscape of head and neck cancer. Nat Rev Cancer 2018, 18:269–282.

8. Zhu L, Wang Y, Li R, Liu A, Zhang X, Zuo C, Xu X: Surgical treatment of early tongue squamous cell carcinoma and patient survival. Oncol Lett 2019, 17:5681–5685.

9. Nemade H, Chaitanya SA, Kumar S, A AK, Rao TS, Rao SL: Oncological outcomes of total glossectomy procedure for advanced tongue cancer: a single-centre experience. Int J Oral Maxillofac Surg 2022, 51:152–158.

10. Katna R, Bhosale B, Sharma R, Singh S, Deshpande A, Kalyani N: Oncological outcomes in patients undergoing major glossectomy for advanced carcinoma of the oral tongue. Ann R Coll Surg Engl 2020, 102:514–518.

11. Chen YH, Chen CH, Chien CY, Su YY, Luo SD, Li SH: JMJD3 suppresses tumor progression in oral tongue squamous cell carcinoma patients receiving surgical resection. PeerJ 2022, 10:e13759.

12. Li Q, Tie Y, Alu A, Ma X, Shi H: Targeted therapy for head and neck cancer: signaling pathways and clinical studies. Signal Transduct Target Ther 2023, 8:31.

13. Zhou Y, Yao Z, Lin Y, Zhang H: From Tyrosine Kinases to Tyrosine Phosphatases: New Therapeutic Targets in Cancers and Beyond. Pharmaceutics 2024, 16.

14. Bou Antoun N, Chioni AM: Dysregulated Signalling Pathways Driving Anticancer Drug Resistance. Int J Mol Sci 2023, 24.

15. Hanke JH, Gardner JP, Dow RL, Changelian PS, Brissette WH, Weringer EJ, Pollok BA, Connelly PA: Discovery of a novel, potent, and Src family-selective tyrosine kinase inhibitor. Study of Lck- and FynT-dependent T cell activation. J Biol Chem 1996, 271:695–701.

16. Olivieri A, Manzione L: Dasatinib: a new step in molecular target therapy. Ann Oncol 2007, 18 Suppl 6:vi42-46.

17. Brooks HD, Glisson BS, Bekele BN, Johnson FM, Ginsberg LE, El-Naggar A, Culotta KS, Takebe N, Wright J, Tran HT, Papadimitrakopoulou VA: Phase 2 study of dasatinib in the treatment of head and neck squamous cell carcinoma. Cancer 2011, 117:2112–2119.

18. Fury MG, Baxi S, Shen R, Kelly KW, Lipson BL, Carlson D, Stambuk H, Haque S, Pfister DG: Phase II study of saracatinib (AZD0530) for patients with recurrent or metastatic head and neck squamous cell carcinoma (HNSCC). Anticancer Res 2011, 31:249–253.

19. Hermida-Prado F, Villaronga M, Granda-Díaz R, Del-Río-Ibisate N, Santos L, Hermosilla MA, Oro P, Allonca E, Agorreta J, Garmendia I, et al: The SRC Inhibitor Dasatinib Induces Stem Cell-Like Properties in Head and Neck Cancer Cells that are Effectively Counteracted by the Mithralog EC-8042. J Clin Med 2019, 8.

20. Pufall MA: Glucocorticoids and Cancer. Adv Exp Med Biol 2015, 872:315–333.

21. Kim KN, LaRiviere M, Macduffie E, White CA, Jordan-Luft MM, Anderson E, Ziegler M, Radcliff JA, Jones J: Use of Glucocorticoids in Patients With Cancer: Potential Benefits, Harms, and Practical Considerations for Clinical Practice. Pract Radiat Oncol 2023, 13:28–40.

22. Wu Z, Doondeea JB, Gholami AM, Janning MC, Lemeer S, Kramer K, Eccles SA, Gollin SM, Grenman R, Walch A, et al: Quantitative chemical proteomics reveals new potential drug targets in head and neck cancer. Mol Cell Proteomics 2011, 10:M111.011635.

23. Emaduddin M, Bicknell DC, Bodmer WF, Feller SM: Cell growth, global phosphotyrosine elevation, and c-Met phosphorylation through Src family kinases in colorectal cancer cells. Proc Natl Acad Sci U S A 2008, 105:2358–2362.

24. Sakai R, Iwamatsu A, Hirano N, Ogawa S, Tanaka T, Mano H, Yazaki Y, Hirai H: A novel signaling molecule, p130, forms stable complexes in vivo with v-Crk and v-Src in a tyrosine phosphorylation-dependent manner. Embo j 1994, 13:3748-3756.

25. Lewitzky M, Simister PC, Feller SM: Beyond ‘furballs’ and ‘dumpling soups’ - towards a molecular architecture of signaling complexes and networks. FEBS Lett 2012, 586:2740–2750.

26. Camacho Leal Mdel P, Sciortino M, Tornillo G, Colombo S, Defilippi P, Cabodi S: p130Cas/BCAR1 scaffold protein in tissue homeostasis and pathogenesis. Gene 2015, 562:1-7.

27. Tikhmyanova N, Little JL, Golemis EA: CAS proteins in normal and pathological cell growth control. Cell Mol Life Sci 2010, 67:1025–1048.

28. Barrett A, Pellet-Many C, Zachary IC, Evans IM, Frankel P: p130Cas: a key signalling node in health and disease. Cell Signal 2013, 25:766–777.

29. Defilippi P, Di Stefano P, Cabodi S: p130Cas: a versatile scaffold in signaling networks. Trends Cell Biol 2006, 16:257–263.

30. Liu D, Peterson ME, Long EO: The adaptor protein Crk controls activation and inhibition of natural killer cells. Immunity 2012, 36:600–611.

31. Mitra SK, Schlaepfer DD: Integrin-regulated FAK-Src signaling in normal and cancer cells. Curr Opin Cell Biol 2006, 18:516–523.

32. Bain J, Plater L, Elliott M, Shpiro N, Hastie CJ, McLauchlan H, Klevernic I, Arthur JS, Alessi DR, Cohen P: The selectivity of protein kinase inhibitors: a further update. Biochem J 2007, 408:297–315.

33. Brandvold KR, Santos SM, Breen ME, Lachacz EJ, Steffey ME, Soellner MB: Exquisitely specific bisubstrate inhibitors of c-Src kinase. ACS Chem Biol 2015, 10:1387–1391.

34. Lee S, Park S, Ryu JS, Kang J, Kim I, Son S, Lee BS, Kim CH, Kim YS: c-Src inhibitor PP2 inhibits head and neck cancer progression through regulation of the epithelial-mesenchymal transition. Exp Biol Med (Maywood) 2023, 248:492–500.

35. Piwnica-Worms H, Saunders KB, Roberts TM, Smith AE, Cheng SH: Tyrosine phosphorylation regulates the biochemical and biological properties of pp60c-src. Cell 1987, 49:75–82.

36. Lindauer M, Hochhaus A: Dasatinib. Recent Results Cancer Res 2018, 212:29–68.

37. Demetri GD, Lo Russo P, MacPherson IR, Wang D, Morgan JA, Brunton VG, Paliwal P, Agrawal S, Voi M, Evans TR: Phase I dose-escalation and pharmacokinetic study of dasatinib in patients with advanced solid tumors. Clin Cancer Res 2009, 15:6232–6240.

38. Mayer BJ, Hirai H, Sakai R: Evidence that SH2 domains promote processive phosphorylation by protein-tyrosine kinases. Curr Biol 1995, 5:296–305.

39. Sakai R, Nakamoto T, Ozawa K, Aizawa S, Hirai H: Characterization of the kinase activity essential for tyrosine phosphorylation of p130Cas in fibroblasts. Oncogene 1997, 14:1419–1426.

40. Manley PW, Cowan-Jacob SW, Buchdunger E, Fabbro D, Fendrich G, Furet P, Meyer T, Zimmermann J: Imatinib: a selective tyrosine kinase inhibitor. Eur J Cancer 2002, 38 Suppl 5:S19–27.

41. Palmer AC, Sorger PK: Combination Cancer Therapy Can Confer Benefit via Patient-to-Patient Variability without Drug Additivity or Synergy. Cell 2017, 171:1678–1691.e1613.

42. Silva JPN, Pinto B, Monteiro L, Silva PMA, Bousbaa H: Combination Therapy as a Promising Way to Fight Oral Cancer. Pharmaceutics 2023, 15.

43. Mackie AE, Ventresca GP, Fuller RW, Bye A: Pharmacokinetics of intravenous fluticasone propionate in healthy subjects. Br J Clin Pharmacol 1996, 41:539–542.

44. Vargas R, Dockhorn RJ, Findlay SR, Korenblat PE, Field EA, Kral KM: Effect of fluticasone propionate aqueous nasal spray versus oral prednisone on the hypothalamic-pituitary-adrenal axis. J Allergy Clin Immunol 1998, 102:191–197.

45. Madamsetty VS, Mohammadinejad R, Uzieliene I, Nabavi N, Dehshahri A, García-Couce J, Tavakol S, Moghassemi S, Dadashzadeh A, Makvandi P, et al: Dexamethasone: Insights into Pharmacological Aspects, Therapeutic Mechanisms, and Delivery Systems. ACS Biomater Sci Eng 2022, 8:1763–1790.

46. Oakley RH, Cidlowski JA: The biology of the glucocorticoid receptor: new signaling mechanisms in health and disease. J Allergy Clin Immunol 2013, 132:1033–1044.

47. Rothenberger NJ, Stabile LP: Hepatocyte Growth Factor/c-Met Signaling in Head and Neck Cancer and Implications for Treatment. Cancers (Basel) 2017, 9.

48. Den Haese GJ, Walworth N, Carr AM, Gould KL: The Wee1 protein kinase regulates T14 phosphorylation of fission yeast Cdc2. Mol Biol Cell 1995, 6:371–385.

49. Katayama K, Fujita N, Tsuruo T: Akt/protein kinase B-dependent phosphorylation and inactivation of WEE1Hu promote cell cycle progression at G2/M transition. Mol Cell Biol 2005, 25:5725–5737.

50. Ge H, Ke J, Xu N, Li H, Gong J, Li X, Song Y, Zhu H, Bai C: Dexamethasone alleviates pemetrexed-induced senescence in Non-Small-Cell Lung Cancer. Food Chem Toxicol 2018, 119:86–97.

51. Starling S: Glucocorticoid-induced bone loss linked with marrow adipocyte senescence. Nat Rev Endocrinol 2023, 19:312.

52. Patki M, McFall T, Rosati R, Huang Y, Malysa A, Polin L, Fielder A, Wilson MR, Lonardo F, Back J, et al: Chronic p27(Kip1) Induction by Dexamethasone Causes Senescence Phenotype and Permanent Cell Cycle Blockade in Lung Adenocarcinoma Cells Over-expressing Glucocorticoid Receptor. Sci Rep 2018, 8:16006.

53. Ge H, Ni S, Wang X, Xu N, Liu Y, Wang X, Wang L, Song D, Song Y, Bai C: Dexamethasone reduces sensitivity to cisplatin by blunting p53-dependent cellular senescence in non-small cell lung cancer. PLoS One 2012, 7:e51821.

54. Srivastava S, Siddiqui S, Singh S, Chowdhury S, Upadhyay V, Sethi A, Kumar Trivedi A: Dexamethasone induces cancer mitigation and irreversible senescence in lung cancer cells via damaging cortical actin and sustained hyperphosphorylation of pRb. Steroids 2023, 198:109269.

55. Kullmann MK, Grubbauer C, Goetsch K, Jäkel H, Podmirseg SR, Trockenbacher A, Ploner C, Cato AC, Weiss C, Kofler R, Hengst L: The p27-Skp2 axis mediates glucocorticoid-induced cell cycle arrest in T-lymphoma cells. Cell Cycle 2013, 12:2625–2635.

56. Levy JMM, Towers CG, Thorburn A: Targeting autophagy in cancer. Nat Rev Cancer 2017, 17:528–542.

57. Grandér D, Kharaziha P, Laane E, Pokrovskaja K, Panaretakis T: Autophagy as the main means of cytotoxicity by glucocorticoids in hematological malignancies. Autophagy 2009, 5:1198–1200.

58. Liu L, Han S, Xiao X, An X, Gladkich J, Hinz U, Hillmer S, Hoppe-Tichy T, Xu Y, Schaefer M, et al: Glucocorticoid-induced microRNA-378 signaling mediates the progression of pancreatic cancer by enhancing autophagy. Cell Death Dis 2022, 13:1052.

59. Tan YQ, Zhang J, Zhou G: Autophagy and its implication in human oral diseases. Autophagy 2017, 13:225–236.

60. Xu J, Su Y, Xu A, Fan F, Mu S, Chen L, Chu Z, Zhang B, Huang H, Zhang J, et al: miR-221/222-Mediated Inhibition of Autophagy Promotes Dexamethasone Resistance in Multiple Myeloma. Mol Ther 2019, 27:559–570.

61. Jiang L, Xu L, Xie J, Li S, Guan Y, Zhang Y, Hou Z, Guo T, Shu X, Wang C, et al: Inhibition of autophagy overcomes glucocorticoid resistance in lymphoid malignant cells. Cancer Biol Ther 2015, 16:466–476.

62. SenthilKumar G, Skiba JH, Kimple RJ: High-throughput quantitative detection of basal autophagy and autophagic flux using image cytometry. Biotechniques 2019, 67:70–73.

63. Werumeus Buning J, Touw DJ, Brummelman P, Dullaart RPF, van den Berg G, van der Klauw MM, Kamp J, Wolffenbuttel BHR, van Beek AP: Pharmacokinetics of oral hydrocortisone - Results and implications from a randomized controlled trial. Metabolism 2017, 71:7–16.

64. Jayne DR, Gaskin G, Rasmussen N, Abramowicz D, Ferrario F, Guillevin L, Mirapeix E, Savage CO, Sinico RA, Stegeman CA, et al: Randomized trial of plasma exchange or high-dosage methylprednisolone as adjunctive therapy for severe renal vasculitis. J Am Soc Nephrol 2007, 18:2180–2188.

65. Leighl NB, Nirmalakumar S, Ezeife DA, Gyawali B: An Arm and a Leg: The Rising Cost of Cancer Drugs and Impact on Access. Am Soc Clin Oncol Educ Book 2021, 41:1–12.

66. Lee YT, Tan YJ, Oon CE: Molecular targeted therapy: Treating cancer with specificity. Eur J Pharmacol 2018, 834:188–196.

67. Vermorken JB, Mesia R, Rivera F, Remenar E, Kawecki A, Rottey S, Erfan J, Zabolotnyy D, Kienzer HR, Cupissol D, et al: Platinum-based chemotherapy plus cetuximab in head and neck cancer. N Engl J Med 2008, 359:1116–1127.

68. De Cecco L, Nicolau M, Giannoccaro M, Daidone MG, Bossi P, Locati L, Licitra L, Canevari S: Head and neck cancer subtypes with biological and clinical relevance: Meta-analysis of gene-expression data. Oncotarget 2015, 6:9627–9642.

69. Morris LGT, Chandramohan R, West L, Zehir A, Chakravarty D, Pfister DG, Wong RJ, Lee NY, Sherman EJ, Baxi SS, et al: The Molecular Landscape of Recurrent and Metastatic Head and Neck Cancers: Insights From a Precision Oncology Sequencing Platform. JAMA Oncol 2017, 3:244–255.

70. Kitamura N, Sento S, Yoshizawa Y, Sasabe E, Kudo Y, Yamamoto T: Current Trends and Future Prospects of Molecular Targeted Therapy in Head and Neck Squamous Cell Carcinoma. Int J Mol Sci 2020, 22.

71. Weiss RA, Vogt PK: 100 years of Rous sarcoma virus. J Exp Med 2011, 208:2351–2355.

72. Li J, Zhang X, Hou Z, Cai S, Guo Y, Sun L, Li A, Li Q, Wang E, Miao Y: P130cas-FAK interaction is essential for YAP-mediated radioresistance of non-small cell lung cancer. Cell Death Dis 2022, 13:783.

73. Pincini A, Tornillo G, Orso F, Sciortino M, Bisaro B, Leal Mdel P, Lembo A, Brizzi MF, Turco E, De Pittà C, et al: Identification of p130Cas/ErbB2-dependent invasive signatures in transformed mammary epithelial cells. Cell Cycle 2013, 12:2409–2422.

74. Cabodi S, Tinnirello A, Di Stefano P, Bisarò B, Ambrosino E, Castellano I, Sapino A, Arisio R, Cavallo F, Forni G, et al: p130Cas as a new regulator of mammary epithelial cell proliferation, survival, and HER2-neu oncogene-dependent breast tumorigenesis. Cancer Res 2006, 66:4672–4680.

75. Cabodi S, Tinnirello A, Bisaro B, Tornillo G, del Pilar Camacho-Leal M, Forni G, Cojoca R, Iezzi M, Amici A, Montani M, et al: p130Cas is an essential transducer element in ErbB2 transformation. Faseb j 2010, 24:3796-3808.

76. Bisaro B, Sciortino M, Colombo S, Camacho Leal MP, Costamagna A, Castellano I, Montemurro F, Rossi V, Valabrega G, Turco E, et al: p130Cas scaffold protein regulates ErbB2 stability by altering breast cancer cell sensitivity to autophagy. Oncotarget 2016, 7:4442–4453.

77. Guo R, Luo J, Chang J, Rekhtman N, Arcila M, Drilon A: MET-dependent solid tumours - molecular diagnosis and targeted therapy. Nat Rev Clin Oncol 2020, 17:569–587.

78. Puccini A, Marín-Ramos NI, Bergamo F, Schirripa M, Lonardi S, Lenz HJ, Loupakis F, Battaglin F: Safety and Tolerability of c-MET Inhibitors in Cancer. Drug Saf 2019, 42:211–233.

79. Choueiri TK, Halabi S, Sanford BL, Hahn O, Michaelson MD, Walsh MK, Feldman DR, Olencki T, Picus J, Small EJ, et al: Cabozantinib Versus Sunitinib As Initial Targeted Therapy for Patients With Metastatic Renal Cell Carcinoma of Poor or Intermediate Risk: The Alliance A031203 CABOSUN Trial. J Clin Oncol 2017, 35:591–597.

80. Hartmann S, Bhola NE, Grandis JR: HGF/Met Signaling in Head and Neck Cancer: Impact on the Tumor Microenvironment. Clin Cancer Res 2016, 22:4005–4013.

81. Sharma A, Menche J, Huang CC, Ort T, Zhou X, Kitsak M, Sahni N, Thibault D, Voung L, Guo F, et al: A disease module in the interactome explains disease heterogeneity, drug response and captures novel pathways and genes in asthma. Hum Mol Genet 2015, 24:3005–3020.

82. Blanquaert F, Pereira RC, Canalis E: Cortisol inhibits hepatocyte growth factor/scatter factor expression and induces c-met transcripts in osteoblasts. Am J Physiol Endocrinol Metab 2000, 278:E509–515.

83. Hoeben A, Martin D, Clement PM, Cools J, Gutkind JS: Role of GRB2-associated binder 1 in epidermal growth factor receptor-induced signaling in head and neck squamous cell carcinoma. Int J Cancer 2013, 132:1042–1050.

84. Kurupi R, Floros KV, Jacob S, Chawla AT, Cai J, Hu B, Puchalapalli M, Coon CM, Khatri R, Crowther GS, et al: Pharmacologic Inhibition of SHP2 Blocks Both PI3K and MEK Signaling in Low-epiregulin HNSCC via GAB1. Cancer Res Commun 2022, 2:1061–1074.

85. Dash S, Hanson S, King B, Nyswaner K, Foss K, Tesi N, Harvey MJB, Navarro-Marchal SA, Woods A, Poradosu E, et al: The SRC family kinase inhibitor NXP900 demonstrates potent antitumor activity in squamous cell carcinomas. J Biol Chem 2024, 300:107615.

86. Hollis RL, Elliott R, Dawson JC, Ilenkovan N, Matthews RM, Stillie LJ, Oswald AJ, Kim H, Llaurado Fernandez M, Churchman M, et al: High throughput screening identifies dasatinib as synergistic with trametinib in low grade serous ovarian carcinoma. Gynecol Oncol 2024, 186:42–52.

87. Borowicz S, Van Scoyk M, Avasarala S, Karuppusamy Rathinam MK, Tauler J, Bikkavilli RK, Winn RA: The soft agar colony formation assay. J Vis Exp 2014:e51998.

88. Uphoff CC, Drexler HG: Comparative PCR analysis for detection of mycoplasma infections in continuous cell lines. In Vitro Cell Dev Biol Anim 2002, 38:79–85.

89. Team RC: R: A Language and Environment for Statistical Computing. Vienna, Austria: R Foundation for Statistical Computing; 2023.

90. Ritz C, Baty F, Streibig JC, Gerhard D: Dose-Response Analysis Using R. PLOS ONE 2015, 10.

91. Vallat R: Pingouin: statistics in Python. Journal of Open Source Software 2018, 3:1026.

92. team Tpd: pandas-dev/pandas: Pandas. Zenodo; 2023.

93. Harris CR, Millman KJ, van der Walt SJ, Gommers R, Virtanen P, Cournapeau D, Wieser E, Taylor J, Berg S, Smith NJ, et al: Array programming with NumPy. Nature 2020, 585:357–362.

94. Caswell TA, Andrade ESd, Lee A, Droettboom M, Hoffmann T, Klymak J, Hunter J, Firing E, Stansby D, Varoquaux N, et al: matplotlib/matplotlib: REL: v3.7.2. Zenodo; 2023.

95. Waskom ML: seaborn: statistical data visualization. Journal of Open Source Software 2021, 6:3021.

